# Nuclear cytoophidia assembly represses transcriptional activity to control skeletal development and homeostasis

**DOI:** 10.1101/2023.12.10.571026

**Authors:** Cheng Xu, Zhixin Wei, Longfei Lv, Xiaoyu Dong, Wenwen Xia, Junqiao Xing, Hongni Liu, Xue Zhao, Yuan Liu, Weihua Wang, Haochen Jiang, Yeli Gong, Cong Liu, Kai Xu, Siyuan Wang, Yoshie Akimoto, Zhangfeng Hu

## Abstract

Compartmentation via filamentation is an evolutionarily conserved subcellular structure that fine-tunes the inherent activity of proteins. Cytoophidia represent a typical class of filamentous structures controlling enzymatic activities. Despite eukaryotic cells containing both cytoplasmic cytoophidia and nuclear cytoophidia, the physiological significance of nuclear cytoophidia is largely unknown. Here we show that nuclear filamentation inhibits the transcriptional activity of Impdh2 required for limb formation and bone resorption. Impdh2 deletion in mouse limb mesenchymal progenitors causes severe skeletal dysplasia by impairing endochondral ossification and chondrocyte differentiation. Additionally, Impdh2 deficiency in myeloid lineages leads to an increased bone mass via impeding osteoclast differentiation. Furthermore, Impdh2 regulates osteoclastic mitochondrial biogenesis and function. We propose that the nuclear compartmentalization of Impdh2 regulates the transcriptional activity during skeletal development and homeostasis.

## INTRODUCTION

Compartmentation, a fundamental character of cellular life, plays a crucial role in many biological processes (Bar-Peled & Kory, 2022, Bhat *et al*, 2021, Tong *et al*, 2022, Zhao & Zhang, 2020). Recently, a new class of compartmentation via filamentation termed cytoophidium or “rod/ring” (RR) has been identified (Aughey & Liu, 2015, Liu, 2010, Liu, 2016, Shen *et al*, 2016). Cytoophidia, which are evolutionarily conserved across prokaryotes (Ingerson-Mahar *et al*, 2010, Zhou *et al*, 2020) and eukaryotes (Ahangari *et al*, 2021, Noree *et al*, 2010, Peng *et al*, 2021), represent intracellular, membraneless and filamentous structures (Liu, 2016, Lynch *et al*, 2020). Furthermore, mutation or deficiency of the proteins that assemble the cytoophidia will cause many diseases (Liu *et al*, 2022, Martin *et al*, 2020, Martin *et al*, 2014, Ruzzo *et al*, 2013). Eukaryotic cells contain cytoplasmic cytoophidia and nuclear cytoophidia (Ahangari, Munoz et al., 2021, Gou *et al*, 2014, Shen, Kassim et al., 2016). Almost all current studies have been focused on the cytoplasmic cytoophidia and suggested that their formation mediates the activity of the cytoophidia-assembled proteins (Aughey & Liu, 2015, Chang *et al*, 2015). Nevertheless, the physiological function and significance of nuclear cytoophidia remain unclear.

Skeletal development and homeostasis are two critical processes at distinct stages for vertebrates, and their abnormalities will trigger various bone-related diseases (Feng & McDonald, 2011, Kovacs *et al*, 2021). During embryogenesis, much of the skeleton, including limbs, originates from the condensation of mesenchymal cells and is formed through endochondral ossification (Kovacs, Chaussain *et al*., 2021, Long & Ornitz, 2013). Although multiple key factors, such as Sox9 (Akiyama *et al*, 2002, Bell *et al*, 1997) and Hox (Desanlis *et al*, 2020, Mallo, 2018, Song *et al*, 2020), are required for skeletal development (Long & Ornitz, 2013, Salhotra *et al*, 2020), their transcriptional regulatory mechanisms are not fully understood. Once skeletal development is formed, the skeleton undergoes dynamic homeostasis throughout life to maintain a healthy bone. Bone resorption, one pivotal process of skeletal homeostasis, is carried out by osteoclasts (Boyle *et al*, 2003). Osteoclast differentiation regulated by various factors requires a large amount of energy supported by mitochondria (Veis & O’Brien, 2023). Nevertheless, the regulatory mechanisms via mitochondria need further elucidation.

Inosine monophosphate dehydrogenase (Impdh), a rate-limited enzyme in the *de novo* synthesis of guanine nucleotides, has been widely studied as an oncogene in many different kinds of cancer (Huang *et al*, 2018, Kofuji *et al*, 2019, Wang *et al*, 2022). Impdh is highly conserved from prokaryotes to humans (Hedstrom, 2009). In mammals, Impdh has two isoforms: Impdh1 and Impdh2 (Senda & Natsumeda, 1994). Impdh-assembled cytoophidia are found in both the cytoplasm and nucleus (Ahangari, Munoz et al., 2021, Chang, Lin et al., 2015). Remarkably, Impdh has been demonstrated as a transcription factor to regulate cell proliferation in *Drosophila* (Kozhevnikova *et al*, 2012). However, so far, whether Impdh also functions as a transcription factor in mammals, and whether the nuclear Impdh-assembled cytoophidia are associated with its transcriptional activity, have not been investigated.

In this study, we identify that nuclear filamentation represses the transcriptional activity of Impdh2 required for limb formation and bone resorption. Mice lacking Impdh2 in limb mesenchymal progenitors lead to limb developmental defects due to the impaired differentiation of mesenchymal cells into chondrocytes. In addition, Impdh2 deletion in mouse myeloid lineages suppresses osteoclast differentiation to induce a high bone mass. Our findings offer new insights into the regulatory mechanism for transcriptional activity by nuclear filamentation during skeletal development and homeostasis.

## RESULTS

### Impdh2 deficiency in mesenchymal progenitors induces skeletal dysplasia

We first examined the expression of the two isoforms of *Impdh*: *Impdh1* and *Impdh2*, during limb development. In contrast to *Impdh1*, *Impdh2* was highly expressed in both the forelimbs and hindlimbs at the early limb formation stages (Supplementary Fig S1A), indicating that Impdh2, rather than Impdh1, functions on limb formation. By the whole-mount *in situ* hybridization, *Impdh2* was predominantly expressed in the craniofacial primordia and the distal region of limb buds at E10.5. Then, its expression was declined overall with the embryo development, but still relatively highly expressed close to phalanx- forming regions where the mesenchymal progenitor cells condensate to form the digits (Fig 1A and Supplementary Fig S1B). Due to the early embryonic lethality in the *Impdh2* global knockout mice (Gu *et al*, 2000), we generated the limb mesenchymal progenitor cells-specific *Impdh2* knockout mice (hereafter referred to as *Impdh2^Prx1-/-^*) by crossing female *Impdh2^f/f^* mice with male *Prx1-Cre* mice. Both mRNA and protein expressions of Impdh2 were dramatically reduced in the limbs of *Impdh2^Prx1-/-^* mice (Fig 1B, 1C, and Supplementary Fig S1C). Strikingly, all the mutant mice exhibited multiple severe skeletal defects after birth (Fig 1D), including: 1) loss of scapula, radius and ulna; 2) shortened humerus, femur, tibia and clavicle; 3) deformed and fused digits; 4) shortened or fused sterna (Fig 1E and 1F). Remarkably, the dysplasia of *Impdh2^Prx1-/-^* mice in the forelimbs was much more severe than those in the hindlimbs. The elongation of the forelimb was almost completely retardant at E12.5 and E14.5, even at the earlier stage E10.5, whereas the hindlimb elongation was only impaired in the *Impdh2^Prx1-/-^* mice (Supplementary Fig S1D and S1E). *Impdh2* deletion did not distort Mendelian segregation (Supplementary Fig S1F). Although *Impdh2^f/f^* and *Impdh2^f/+^*; *Prx1*- *Cre* littermates were viable, and fertile with normal skeletal phenotypes, only about 25% of *Impdh2^Prx1-/-^* mice survived to adulthood (Fig 1G). Taken together, Impdh2 is essential for limb development.

**Figure 1.**
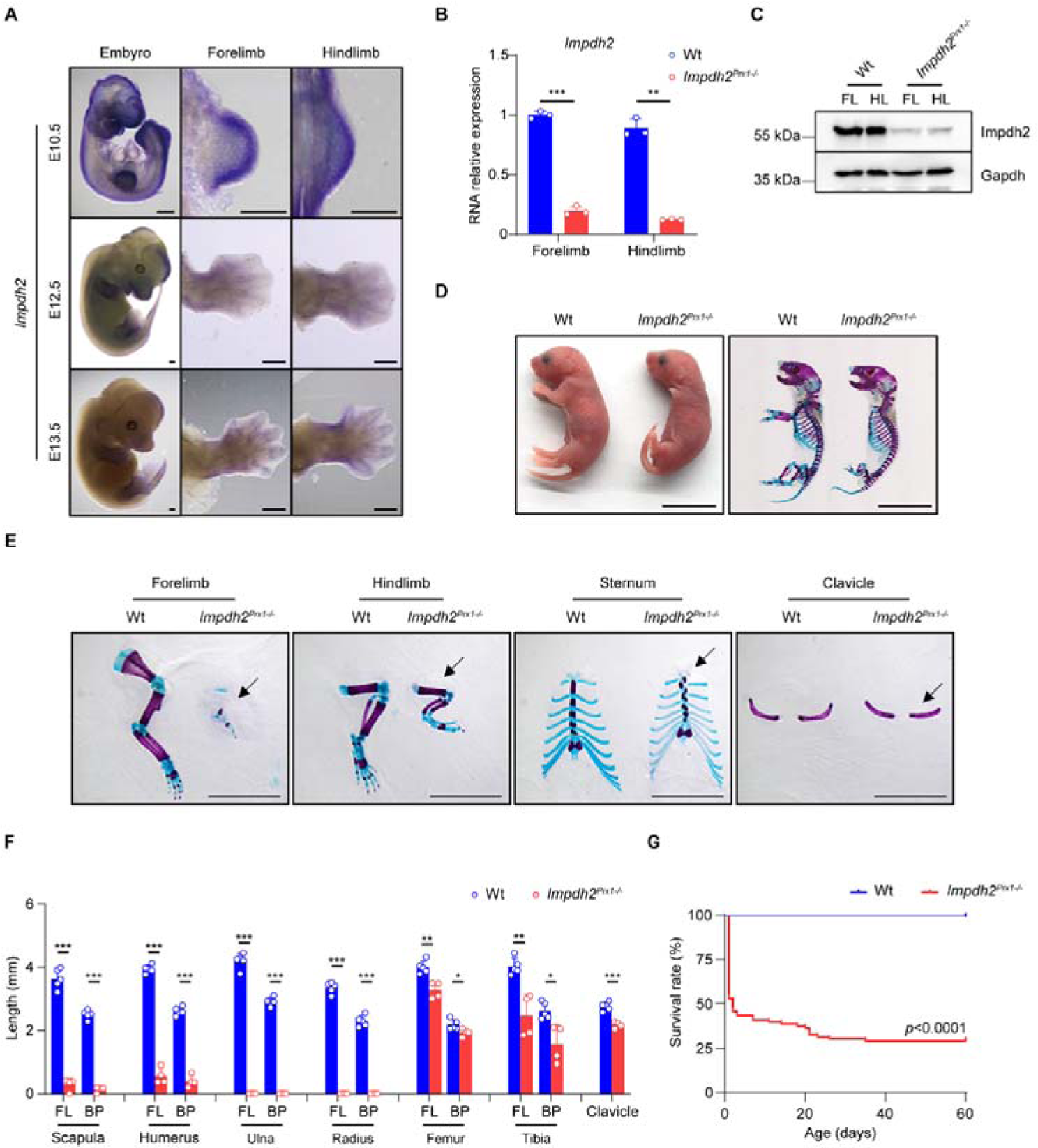
*Impdh2^Prx1-/-^* mice display multiple skeletal abnormalities. A. Whole-mount *in situ* hybridization for *Impdh2* in the mouse embryo (left), forelimbs (middle) and hindlimbs (right) at the indicated stages. Scale bar, 400 µm. B, C. Quantitative qPCR (B) and immunoblot analysis (C). Total RNAs or proteins extracted from the forelimbs and hindlimbs of Wt and *Impdh2^Prx1-/-^* embryos at E12.5 were performed to qPCR (B) and immunoblot analysis (C) for Impdh2 expression. FL: forelimb; HL: hindlimb. Experiments were repeated three times. D. Gross appearance (left) and the skeleton stained with alizarin red and alcian blue (right) from P0 Wt and *Impdh2^Prx1-/-^*mice. Scale bar, 1 cm. E. Forelimb, hindlimb, sternum and clavicle from P0 Wt and *Impdh2^Prx1-/-^*mice. Note that the defects were present in *Impdh2^Prx1-/-^* mice (arrow). Scale bar, 0.5 cm. F. The length of the skeleton from E. Wt, n=5; *Impdh2^Prx1-/-^*, n=4. FL: full length; BP: bone part. G. The survival rate of Wt and *Impdh2^Prx1-/-^* mice after birth. Wt, n=26; *Impdh2^Prx1-/-^*, n=83. The survival time for each mouse was recorded. Kaplan- Meier curves were generated, and datasets were compared (log-rank test) using GraphPad Prism *version 8.0*. In (B, F), results were expressed as mean ± s.d. \**p*<0.05, \*\**p*<0.01, \*\*\**p*<0.001, versus Wt, Student’s *t*-test. See also Supplementary Figure S1.

### Impdh2 deletion attenuates POC and chondrocyte formation

Given that limb elongation was impaired in *Impdh2^Prx1-/-^* mice, we investigated whether the formation of chondrocytes, derived from limb mesenchymal progenitor cell condensation, was impaired. Alcian blue staining was performed to analyze the primary ossification center (POC) formation of the femur, sternum, and humerus at P0. The POC lengths were significantly shorter in *Impdh2^Prx1-/-^*mice (Fig 2A). H&E staining revealed that the heights of the resting zone (RZ), proliferative zone (PZ), and hypertrophic zone (HZ) of the growth plate were significantly decreased in the proximal humerus of *Impdh2^Prx1-/-^*mice (Fig 2B). In WT, the chondrocytes in PZ were discoid and retained a regular arrangement parallel to the long axis of the bone, but stacks of discoid cells were dramatically reduced in the mutant mice. Furthermore, in contrast to the HZ of Wt mice which formed the highly organized and hypertrophied, the chondrocyte organization was messy, and the cell density was significantly decreased in *Impdh2^Prx1-/-^* mice (Fig 2C).

**Figure 2.**
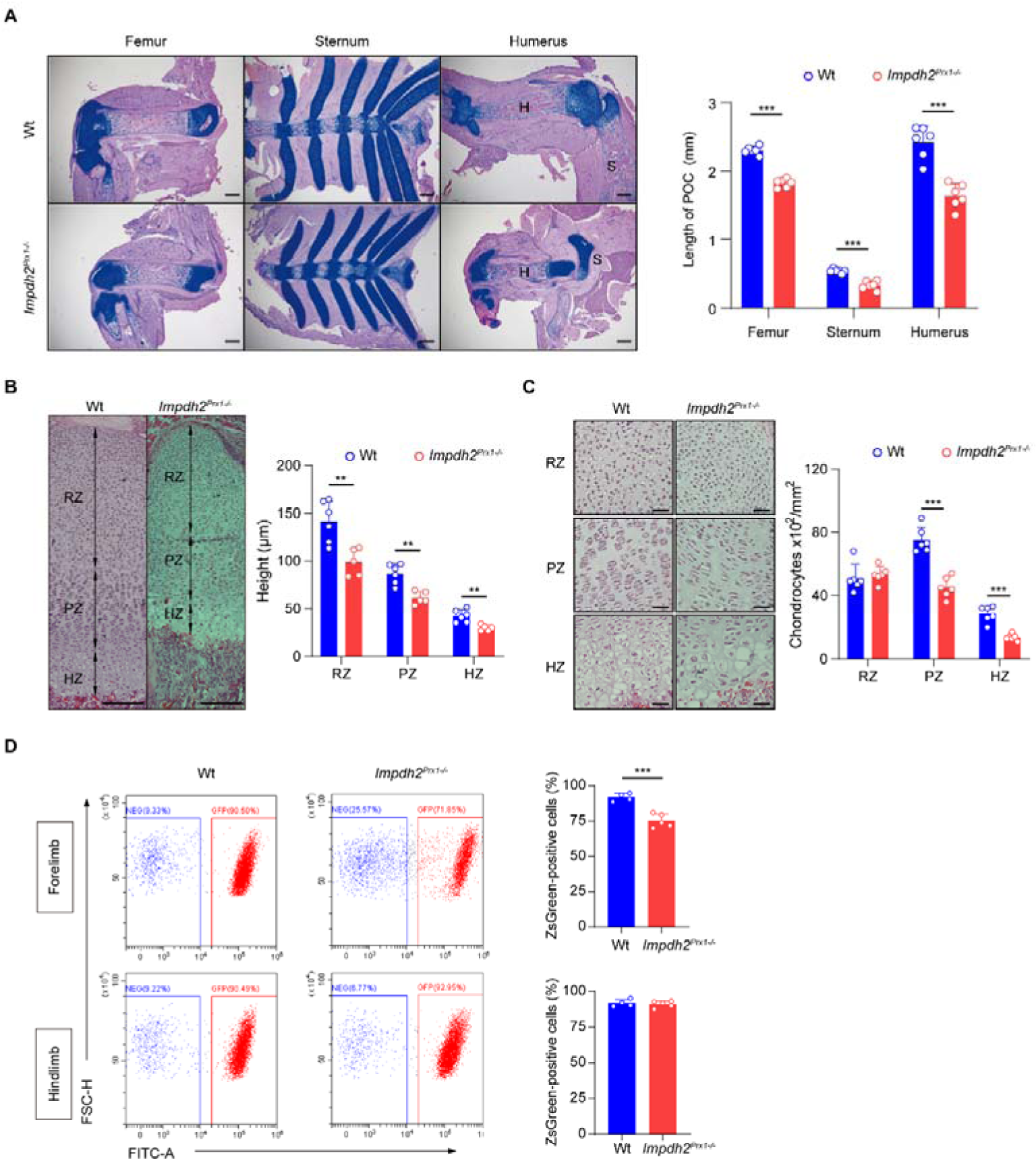
Loss of Impdh2 reduces endochondral ossification and chondrocyte formation. A. Alcian blue staining images (left) and length (right) of POCs in the femur, sternum and humerus from P0 Wt and *Impdh2^Prx1-/-^*mice. S, scapula; H, humerus. n=6/group. Scale bar, 400 µm. B. Hematoxylin staining images (left) and length (right) of the growth plate in proximal humerus from P0 Wt and *Impdh2^Prx1-/-^* mice. RZ, resting zone; PZ, proliferative zone; HZ, hypertrophic zone. Wt, n=6; *Impdh2^Prx1-/-^*, n=5. Scale bar, 50 µm. C. Representative images (left) and quantitation of chondrocyte density (right) of the RZ, PZ and HZ in the humeral growth plate of Wt and *Impdh2^Prx1-/-^* mice. n=6/group. Scale bar, 20 µm. D. Flowcytometry image (left) and quantification (right) of Rosa26-ZsGreen- positive cells of the forelimbs and hindlimbs extracted from Wt and *Impdh2^Prx1-/-^* embryos at E12.5. Wt, n=4; *Impdh2^Prx1-/-^*, n=5. In (A-D), results were expressed as mean ± s.d. \*\**p*<0.01, \*\*\**p*<0.001, versus Wt, Student’s *t*-test.

Considering that the heights of RZ, PZ, and HZ were reduced, we next investigated whether the population of chondrocyte progenitor cells, i.e., the limb mesenchymal cells was affected. The limb mesenchymal cells of Wt and *Impdh2^Prx1-/-^* mice were genetically labeled using the reporter *Rosa26*- *ZsGreen* mice to examine the limb mesenchymal cell population at E12.5. Flow cytometry analysis revealed that the ZsGreen-positive cell population was significantly decreased in the forelimbs of *Impdh2^Prx1-/-^* mice, whereas no difference was found in the hindlimbs (Fig 2D). This difference between forelimbs and hindlimbs might partially explain why forelimb defects were more severe than hindlimb in *Impdh2^Prx1-/-^*mice.

### Impdh2 controls chondrogenesis through transcriptionally regulating bone development-related gene expression

To investigate the potential mechanisms that impdh2 regulates limb formation, we performed high throughput transcriptome sequencing (RNA-seq) using the forelimb buds of control and *Impdh2^Prx1-/-^* mice at E12.5. Two biological replicates were performed to evaluate the reproducibility of RNA-seq data in this study. The RNA-seq data revealed that compared to Wt mice, 841 genes were downregulated, while 723 genes were upregulated in *Impdh2^Prx1-/-^* mice with the following screening conditions: DE, |log2FoldChange|>0.5, *p*<0.05 (Fig 3A). Based on gene ontology (GO) analysis with the downregulated genes, the top 10 biological process terms were highly involved in chondrogenesis and skeletogenesis, including chondrocyte differentiation, cartilage development, skeletal system development and limb morphogenesis (Fig 3B). We further extracted these gene expression values from the RNA- seq data and confirmed that 55 chondrogenic and skeletogenic genes were dramatically suppressed in *Impdh2^Prx1-/-^* mice, such as *Sox9*, *Ihh* and *Aggrecan* (*Acan*) (Fig 3C). Quantitative PCR was performed to further confirm that the expressions of *Sox9* and *Col2a1* were dramatically suppressed by Impdh2 deficiency during limb formation (Fig 3D). Furthermore, the expression patterns of *Sox9* and *Col2a1*, especially in the forelimbs, were strikingly impaired in *Impdh2^Prx1-/-^*mice by the whole-mount *in situ* hybridization detection (Fig 3E). Then, micromass cultures were performed and the results showed that chondrogenic nodule formation and marker gene expressions, such as *Sox9*, *Col2a1* and *Acan*, were significantly decreased in *Impdh2^Prx1-/-^* mice (Fig 3F, 3G, and 3H). Additionally, it’s well known that the transcription factors, *Hox* genes are essential for limb morphology and skeleton formation during embryogenesis. In particular, Hoxa11 and Hoxd11 are indispensable for radius and ulna formation (Davis *et al*, 1995, Song, Pineault et al., 2020, Wellik & Capecchi, 2003), while Hoxa13 and Hoxd13 are mandatory for digits development (Desanlis, Kherdjemil et al., 2020, Fromental-Ramain *et al*, 1996). Due to these similar phenotypes with the *Impdh2^Prx1-/-^* mice, we hypothesized that *Impdh2* deficiency also causes mice limb abnormality by controlling the *Hox* gene transcriptionally expression.

**Figure 3.**
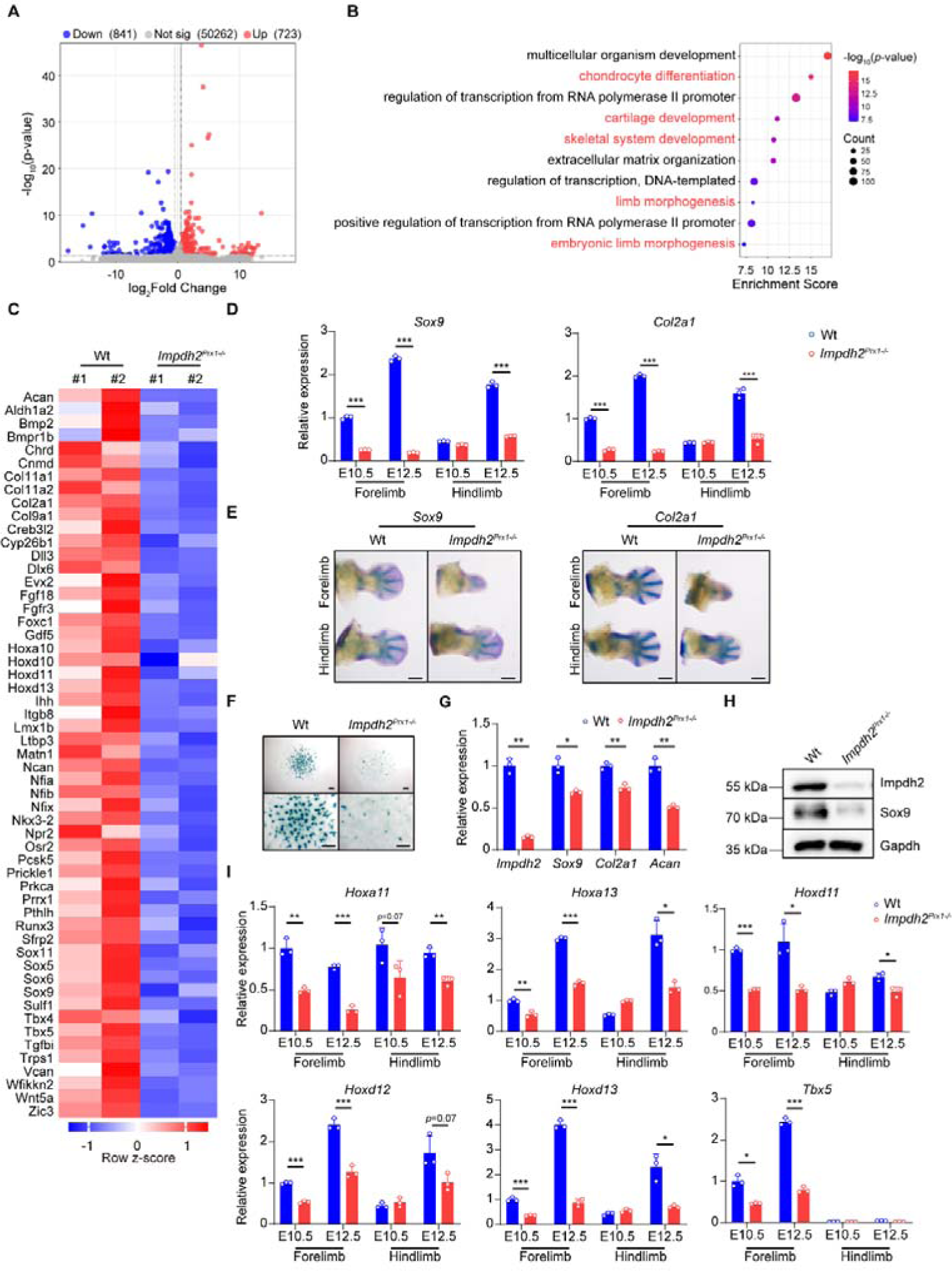
Deletion of Impdh2 suppresses chondrocyte differentiation. A. Volcano plot for comparison of RNA-seq data of the forelimbs from Wt and *Impdh2^Prx1-/-^* embryos at E12.5. Nondifferent genes are shown in grey, while downregulated genes (log2FoldChange<-0.5, *p*<0.05) and upregulated genes (log2FoldChange>0.5, *p*<0.05) are marked with blue and red color, respectively. B. Biology process (BP) analysis of downregulated genes from (A). C. Heatmaps of mRNA expression of the downregulated genes involved in chondrocyte differentiation, skeletal system development, cartilage development and limb morphogenesis by *Impdh2* deficient. Row z-score of FPKMs of genes was shown in the heatmap. #1, replicate 1. #2, replicate 2. D. Quantitative qPCR analysis of *Sox9* (left) and *Col2a1* (right) expression in the forelimbs and hindlimbs of Wt and *Impdh2^Prx1-/-^* embryo at E10.5 and E12.5, respectively. Experiments were repeated three times. E. Whole-mount *in situ* hybridization for *Sox9* and *Col2a1* in the forelimbs and hindlimbs of Wt and *Impdh2^Prx1-/-^* embryo at E12.5. Scale bar, 400 µm. F. Alcian blue staining images of micromass culture from the Wt and *Impdh2^Prx1-/-^* forelimbs. Primary mesenchymal progenitor cells isolated from the Wt and *Impdh2^Prx1-/-^* forelimbs at E12.5 were cultured and stimulated with ascorbic acid and β-glycerol phosphate for 6 days. Scale bar, 400 µm. G. Quantitative qPCR analysis of *Impdh2, Sox9*, *Col2a1* and *Acan* expression from F. H. Western blot analysis of Impdh2 and Sox9 expression from F. Experiments were repeated three times. I. Quantitative qPCR analysis of *Hoxa11*, *Hoxa13*, *Hoxd11*, *Hoxd12*, *Hoxd13* and *Tbx5* expression in the forelimbs and hindlimbs of Wt and *Impdh2^Prx1-/-^* embryos at E10.5 and E12.5. Experiments were repeated three times. In (D, G, I), results were expressed as mean ± s.d. \**p*<0.05, \*\**p*<0.01, \*\*\**p*<0.001, versus Wt, Student’s *t*-test. See also Supplementary Figure S2.

Indeed, the expression of *Hox* genes, such as *Hoxa10*, *Hoxd10*, *Hoxd11* and *Hoxd13*, was decreased in our RNA-seq data (Fig 3C). The expression of *Hox* genes, such as *Hoxa11*, *Hoxa13*, *Hoxd11*, *Hoxd12*, *Hoxd13* and the target gene *Tbx5* required for the forelimbs, not hindlimbs formation, were strongly impaired during limb formation (Fig 3I and Supplementary Fig S2). Collectively, these results indicated that *Impdh2* deletion inhibits the differentiation of mesenchymal cells into chondrocytes by regulating the transcriptional expression of bone development-related genes.

### *Impdh2^Prx1-/-^* adult mice display abnormal skeletons

Given that approximately 25% of *Impdh2^Prx1-/-^* mice successfully survived to adulthood (Fig 1G), we further investigated the bone phenotypes of these adult mice. The body weights were significantly decreased in the mutant mice (Fig 4A). In line with the previous observation at P0, the *Impdh2^Prx1-/-^* adult mice also displayed multiple skeletal deformities, such as absent radius and ulna; shortened humerus, femur and tibia; fused and shortened digits (Fig 4B, 4C, and 4D). We next tried to extract bone marrow mesenchymal stromal cells (BMSCs) to perform osteogenic experiments. However, the cells from *Impdh2^Prx1-/-^* mice did not proliferate (Fig 4E), indicating that *Impdh2* absence suppresses BMSCs proliferation. Nevertheless, no significant difference was found in the bone mass of lumbar vertebrae between 12-week-old male Wt and *Impdh2^Prx1-/-^* mice (Fig 4F). Considering the possibility that the Cre driven by the *Prx1* promoter mainly functions in long bones, the bone mass of the femur was also analyzed by µCT analysis. In contrast with the Wt mice, both the tissue volume and bone volume were significantly reduced in *Impdh2^Prx1-/-^*mice, whereas the BV/TV and the thickness of trabecular bone were not altered. The space of trabecular bone was increased, while the trabecular number tended to decrease (Fig 4G and 4H). Remarkably, the morphology of the distal femur was abnormal in the mutant mice (Fig 4G). Additionally, *Impdh2* absence did not affect the cortical thickness (Fig 4I and 4J).

**Figure 4.**
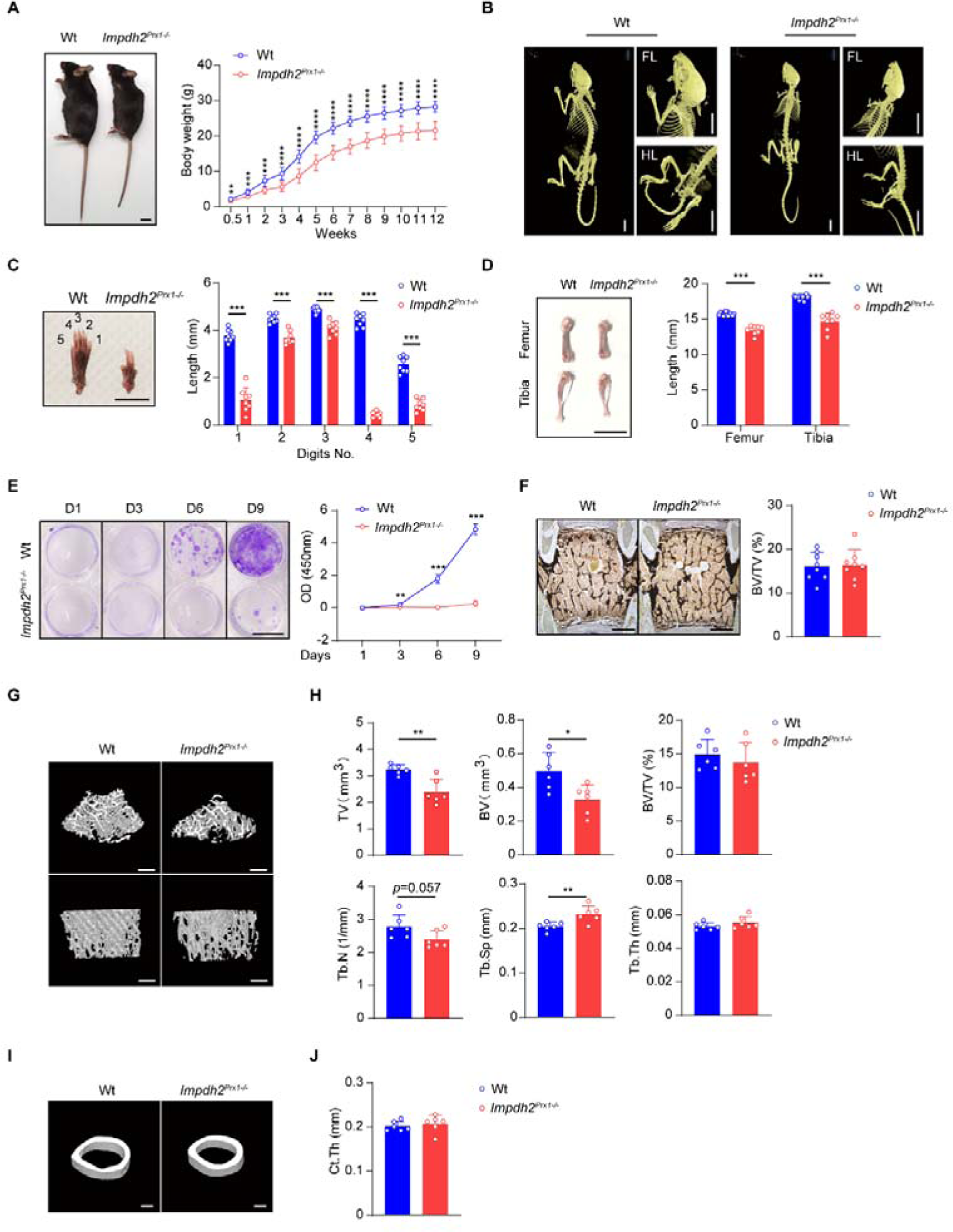
Loss of Impdh2 causes multiple skeletal defects in adult mice. A. Gross appearance of 12-week-old male Wt and *Impdh2^Prx1-/-^* mice (left) and body weight changes (right) in male mice. Wt, n=12; *Impdh2^Prx1-/-^*, n=10. Scale bar, 1 cm. B. Representative µCT images of male Wt and *Impdh2^Prx1-/-^*mice. Note that *Impdh2^Prx1-/-^* mice showed skeletal defects in the forelimbs (FL) and hindlimbs (HL). Scale bar, 1 cm. C. Gross appearance of digits (left) and the length (right) in the indicated digit numbers from 12-week-old male Wt and *Impdh2^Prx1-/-^* mice. n=8/group. Scale bar, 1 cm. D. Gross appearance and length of tibia and femur from 12-week-old male Wt and *Impdh2^Prx1-/-^* mice. Wt, n=18; *Impdh2^Prx1-/-^*, n=8. Scale bar, 1 cm. E. Crystal violet staining and quantitative analysis of bone marrow stromal cells isolated from 12-week-old male Wt and *Impdh2^Prx1-/-^*mice. n=4/group. Scale bar, 1 cm. F. Histological analysis of the vertebrae in 12-week-old male Wt and *Impdh2^Prx1-/-^* mice. n=8/group. Scale bar, 100 µm. G, H. µCT images (G) and bone morphometric analysis (H) of trabecular bone of the distal femurs isolated from 12-week-old male Wt and *Impdh2^Prx1-/-^* mice. TV, tissue volume; BV, bone volume; BV/TV, bone volume/tissue volume; Tb.N, trabecular number; Tb.Sp, trabecular separation; Tb.Th, trabecular thickness. n=6/group. Scale bar, 500 µm. I, J. µCT images (I) and bone morphometric analysis (J) of cortical bone of the mid-shaft femurs isolated from 12-week-old male Wt and *Impdh2^Prx1-/-^* mice. Ct.Th, cortical thickness. n=6/group. Scale bar, 500 µm. In (A-E, H), results were expressed as mean ± s.d. \**p*<0.05, \*\**p*<0.01, \*\*\**p*<0.001, \*\*\*\**p*<0.0001, versus Wt, Student’s *t*-test.

### *Impdh2* deficiency in osteoclasts induces high bone mass by suppressing osteoclastogenesis

Since Impdh2 is indispensable for bone development, we next investigated whether Impdh2 functions the bone resorption, one critical process for bone homeostasis. Bone resorption is mediated by osteoclasts. Excessive osteoclast activity disrupts bone homeostasis and causes many bone diseases, such as inflammatory arthritis and osteoporosis. First, *Impdh2* expression was examined in the various tissues including bone, and the bone- related cell lineages, such as chondrocytes, osteoblasts and osteoclasts. The high expression in osteoclasts indicated that Impdh2 might play a role in osteoclastogenesis (Supplementary Fig S3A and S3B). To investigate the function of Impdh2 on bone resorption, we generated *Impdh2* deletion in myeloid lineage cells, i.e., osteoclast precursors, with *LysM*-*Cre* mice (hereafter referred to as *Impdh2^LysM-/-^*). After RNAKL stimulation, *Impdh2* mRNA and protein expression were first increased at day 1 and then dropped off to the initial level during osteoclast differentiation in Wt mice, whereas they were significantly decreased in *Impdh2^LysM-/-^* mice (Fig 5A and 5B). Notably, the bone mass of lumbar vertebrae was increased in the mutant mice (Fig 5C). Bone histomorphometric analysis showed that the osteoclast surface and number were significantly reduced in *Impdh2^LysM-/-^* mice (Fig 5D). In line with the observations, a high bone mass was also found in the femur of *Impdh2^LysM-/-^* mice by µCT analysis. Meanwhile, the trabecular number was increased without altering the trabecular space and trabecular thickness (Fig 5E and 5F). The cortical thickness was not affected (Supplementary Fig S4A and S4B). To further explore the function of *Impdh2* on osteoclastogenesis *in vitro*, bone marrow cells from Wt and *Impdh2^LysM-/-^* mice were cultured to induce osteoclastic differentiation by M-SCF and RANKL stimulation. The number of TRAP-positive multinucleated osteoclasts was dramatically decreased in response to RANKL in *Impdh2^LysM-/-^* mice (Fig 5G). Furthermore, the expression of marker genes, *Nfatc1*, *Ctsk*, *Calcr* and *Acp5*, as well as the fusion-related genes, *Dcstamp* and *Atp6v0d2*, was significantly decreased (Fig 5H). Moreover, the decreased expression of osteoclastic transcript factors Nfact1, Blimp1 and c-Fos, was further confirmed in the mutant mice by western blot (Fig 5I). Taken together, these results clearly showed that Impdh2 absence causes high bone mass via repressing osteoclast differentiation.

**Figure 5.**
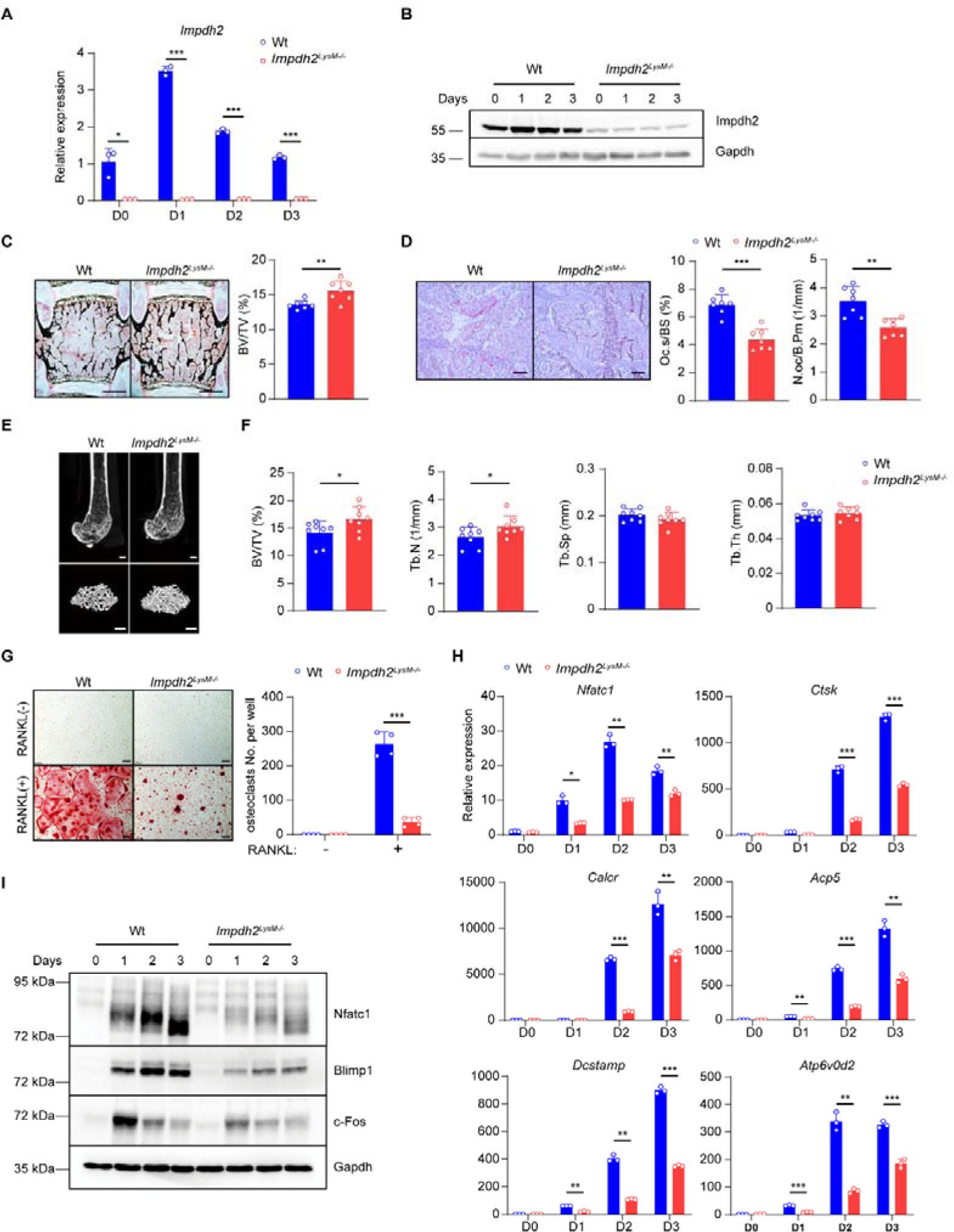
Deletion of Impdh2 in osteoclasts induces high bone mass via suppressing osteoclastogenesis. A, B. Quantitative qPCR (A) and immunoblot analysis (B) of Impdh2 expression during osteoclastogenesis in Wt and *Impdh2^LysM-/-^*osteoclast cultures induced by RANKL. Experiments were repeated three times. C. Histological analysis of the vertebrae in 12-week-old male Wt and *Impdh2^LysM-/-^* mice. n=7/group. Scale bar, 100 µm. D. Histomorphometric analysis of the vertebrae in 12-week-old male Wt and *Impdh2^LysM-/-^* mice. Oc.S/BS, osteoclast surface/bone surface; N.Oc/B.Pm, osteoclast number/bone perimeter. n=7/group. Scale bar, 50 µm. E, F. µCT images (E) and bone morphometric analysis (F) of trabecular bone of the distal femurs isolated from 12-week-old male Wt and *Impdh2^LysM-/-^* mice. n=8/group. Scale bar, 500 µm. G. Osteoclast differentiation derived from BMMs of Wt and *Impdh2^LysM-/-^* mice stimulated with RANKL. TRAP staining was performed and the number of TRAP-positive multinuclear osteoclasts (≥3 nuclei/cell) per well was calculated. TRAP-positive cells appear red in the photographs. Experiments were repeated three times. Scale bar, 100 µm. H. Quantitative qPCR analysis of mRNA expression of the indicated genes during osteoclastogenesis in Wt and *Impdh2^LysM-/-^* osteoclast cultures induced by RANKL at the indicated times. I. Immunoblot analysis of the expression of Nfatc1, Blimp1 and c-Fos in Wt and *Impdh2^LysM-/-^* osteoclast cultures induced by RANKL at the indicated times. Experiments were repeated three times. In (A, C, D, F-H), results were expressed as mean ± s.d. \**p*<0.05, \*\**p*<0.01, \*\*\**p*<0.001, versus Wt, Student’s *t*-test. See also Supplementary Figures S3 and S4.

### Impdh2 deletion represses osteoclastic mitochondrial biogenesis and function and alleviates OVX-induced bone loss

To further explore the mechanisms by which Impdh2 functions on osteoclast differentiation, we also performed high throughput transcriptome sequencing using the WT and *Impdh2^LysM-/-^* osteoclasts with/without RANKL stimulation to identify genes transcriptionally regulated by Impdh2. In the study, two batches of RNA-seq experiments were performed for each condition. To eliminate the differences in the degree of osteoclast differentiation between batches, we first screened the differentially expressed genes in each batch with the following conditions: DE, |log2FoldChange|>0.3, *p*<0.05, and then performed GO analysis with the shared expression genes. After RNAKL stimulation, a total of 613 downregulated and 392 upregulated genes by Impdh2 deficiency were identified (Supplementary Fig S5A). RNA-seq-based expression heatmap of osteoclastic transcription factors and marker genes, such as *Nfact1*, *Ctsk*, *Calcr*, *Acp5*, *Dcstamp* and *Atp6v0d2*, was strongly suppressed in osteoclasts of the *Impdh2^LysM-/-^*mice (Fig 6A). GO analysis revealed that the significantly downregulated genes by *Impdh2* deficiency were highly involved in mitochondrial structures and functions, including oxidative phosphorylation (OXPHOS) (Fig 6B). The expression value of these genes (FPKM) involved in mitochondrion components and OXPHOS pathway from our RNA-seq data was extracted and confirmed that 93 mitochondrion-related genes and 28 OXPHOS-related genes were suppressed in the *Impdh2^LysM-/-^* osteoclasts (Fig 6C and Supplementary Fig S5B). Since osteoclastogenesis is a highly energy-consuming process, the inhibition of osteoclast differentiation by *Impdh2* deficiency might be due to insufficient energy supported by mitochondria. qPCR analysis revealed that the expression of OXPHOS complex components, such as complex I (*Ndufb6*), complex III (*Uqcrq*, *Uqcr10*, *Cyc1*), complex IV (*Cox6a1*) and complex V (*Atp6v1e1*, *Atp5g3*, *Atp6v1b2*), was significantly decreased in the *Impdh2^LysM-/-^* osteoclasts (Fig 6D). Western blot analysis further confirmed that the OXPHOS complexes were repressed by Impdh2 deficiency in RANKL-induced osteoclastogenesis (Fig 6E). Furthermore, the mitochondrial DNA content was also decreased in the *Impdh2^LysM-/-^* osteoclasts (Fig 6F).

**Figure 6.**
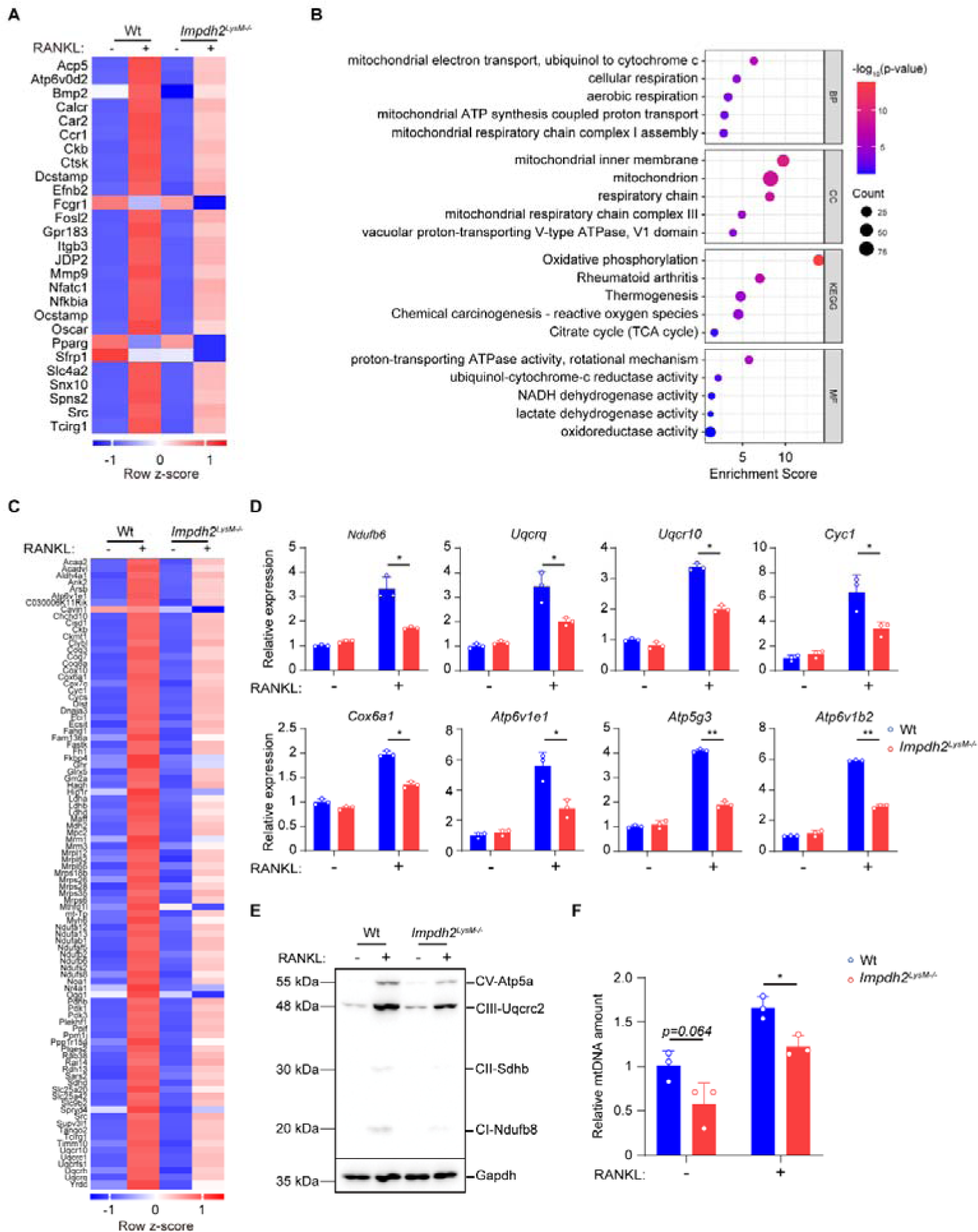
Impdh2 regulates mitochondrial biogenesis and function in osteoclasts. A. Heatmaps of RANKL-induced osteoclast marker genes and transcription factors regulated by Impdh2 deficiency. Row z-score of FPKMs of genes was shown in the heatmap. B. Gene ontology analysis of RANKL-inducible downregulated genes by Impdh2 deficiency. BP, biological process; CC, cellular component; MF, molecular function; KEGG, Kyoto Encyclopedia of Genes and Genomes. C. Heatmaps of mRNA expression of mitochondria-related downregulated genes by Impdh2 deficiency. Row z-score of FPKMs of genes was shown in the heatmap. D. Quantitative qPCR analysis of OXPHOS-related gene expression during RANKL-induced osteoclastogenesis in Wt and *Impdh2^LysM-/-^*mice. Experiments were repeated three times. E. Immunoblot analysis of the OXPHOS expression during RANKL-induced osteoclastogenesis in Wt and *Impdh2^LysM-/-^* mice. F. Relative mitochondrial DNA (mtDNA) amount analysis. The total DNA was isolated from the Wt and *Impdh2^LysM-/-^* osteoclasts stimulated by RANKL for 4 days, and the amount of mitochondrial/chromosol DNA was measured by qPCR analysis using the NADH dehydrogenase subunit 6 (*Nd6*) gene and *Sox9* gene loci. In (D, F), results were expressed as mean ± s.d. \**p*<0.05, \*\**p*<0.01, versus Wt, Student’s *t*-test. See also Supplementary Figures S5 and S6.

Considering that osteoporosis, one of the most severe bone loss diseases, commonly occurs in postmenopausal women due to estrogen deficiency, we established the bilateral ovariectomy (OVX) mouse model to investigate the significance of Impdh2 in pathological bone loss. To evaluate the success of the OVX-induced osteoporosis model, the uterine weights were measured 5 weeks after surgery. Compared with the sham group, the uterine weights were significantly decreased by approximately 70% in OVX groups (Supplementary Fig S6A), suggesting that the estrogen was comparably and effectively deprived in both Wt and *Impdh2^LysM-/-^* mice. As expected, in Wt mice, compared with the sham group, BV/TV and trabecular number were decreased, while trabecular space was increased after OVX surgery (Supplementary Fig S6B and S6C). In the sham group, the BV/TV and trabecular number of *Impdh2^LysM-/-^* female mice were increased compared with Wt sham group mice, consistent with the results of male mice (Fig 5E and 5F). Furthermore, in the OVX group, BV/TV and the trabecular number of *Impdh2^LysM-/-^*mice were significantly increased compared with Wt mice (Supplementary Fig S6B and S6C). Loss of *Impdh2* appeared to not alter the cortical bone thickness after OVX surgery (Supplementary Fig S6D).

### Nuclear cytoophidia assembly suppresses Impdh2 transcriptional activity

Mycophenolic acid (MPA), as a potent uncompetitive IMPDH inhibitor, is approved by the U.S. Food and Drug Administration (FDA) to prevent the rejection of organ transplantation. Thus, we investigated whether MPA inhibits chondrogenesis and osteoclastogenesis. First, in line with the previous results from *Impdh2^Prx1-/-^* mice, Impdh2 inhibition by MPA treatment dramatically decreased the chondrogenic nodule formation (Fig 7A). Meanwhile, the transcriptional expression of the chondrogenic marker genes*, Sox9*, *Col2a1* and *Acan*, was significantly decreased (Fig 7B and 7C). Unexpectedly, the Impdh2 protein expression level, not the mRNA level, was strongly increased after MPA treatment (Fig 7B and 7C). Impdh functions as a transcription factor in the nucleus (Kozhevnikova, van der Knaap et al., 2012). Interestingly, Impdh2 expression was also strongly increased in the nucleus (Fig 7D), which appeared to contradict the previous conclusion that the transcription factor Impdh2 plays a positive role in chondrogenesis. Given that Impdh2 assembles to form a “rods/rings” (RR) structure, i.e., Impdh2-cytoophidia, by MPA stimulation (Keppeke *et al*, 2018, Schiavon *et al*, 2018), we investigated whether nuclear Impdh2-cytoophidia formation mediates its transcriptional activity during chondrogenesis. The nuclear Impdh2-cytoophidia were observed in the limb mesenchymal progenitors *in vivo*, indicating that nuclear Impdh2-cytoophidia might function during limb formation (Supplementary Fig S7A). Additionally, tons of Impdh2-cytoophidia were assembled by MPA treatment in the limb mesenchymal cells *in vitro* (Supplementary Fig S7B). Furthermore, the nuclear Impdh2-cytoophidia were significantly accumulated (Fig 7E and 7F). These collective results suggested that despite of the increased expression of Impdh2 in the nucleus by MPA treatment, the nuclear Impdh2-cytoophidia assembly inhibits Impdh2 transcriptional activity during the chondrocyte differentiation.

**Figure 7.**
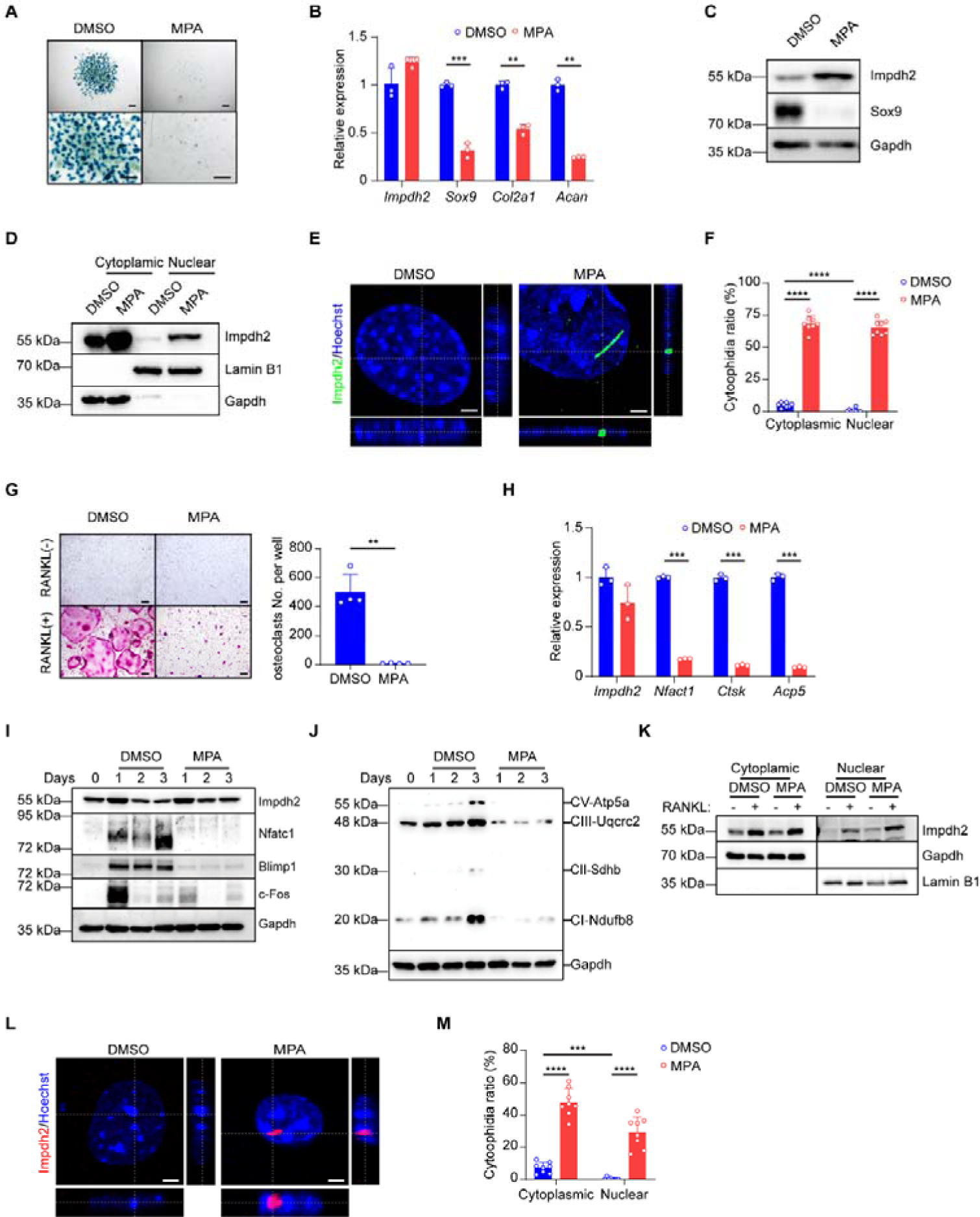
Impdh2-cytoophidia formation mediates chondrogenesis and osteoclastogenesis. A. Alcian blue staining images of micromass culture treated with 1 µM MPA. Primary mesenchymal progenitor cells isolated from Wt limbs at E12.5 were cultured for 6 days supplied with/without 1 µM MPA. Scale bar, 400 µm. B. Quantitative qPCR analysis of *Impdh2, Sox9*, *Col2a1* and *Acan* expression after DMSO or 1 µM MPA treatment. C. Western blot analysis of Impdh2 and Sox9 expression after 1 µM MPA treatment. Experiments were repeated three times. D. Western blot analysis of Impdh2 distribution in primary mesenchymal progenitor cells treated with/without 1 µM MPA. Experiments were repeated three times. E. 3D-projection of images of the nuclear Impdh2-cytoophidia (green) in primary mesenchymal progenitor cells after ascorbic acid and β-glycerol phosphate stimulation for 6 days. Scale bar, 3 µm. F. The ratio of Impdh2-cytoophidia in the cytoplasm and nucleus. Eight random microscope fields of cells were counted. G. Osteoclast differentiation from BMMs derived from Wt mice stimulated with/without 1 µM MPA. TRAP staining was performed and the number of TRAP-positive multinuclear osteoclasts (≥3 nuclei/cell) per well was calculated. Experiments were repeated three times. Scale bar, 100 µm. H. Quantitative qPCR analysis of *Impdh2, Nfatc1*, *Ctsk* and *Acp5* expression during RANKL-induced osteoclastogenesis treated with/without 1 µM MPA. I. Immunoblot analysis of the expression of Impdh2, Nfatc1, Blimp1 and c-Fos in osteoclasts treated with/without 1 µM MPA at the indicated times. J. Immunoblot analysis of the OXPHOS complexes expression during RANKL-induced osteoclastogenesis treated with/without 1 µM MPA at the indicated times. K. Western blot analysis of Impdh2 distribution during RANKL-induced osteoclastogenesis treated with/without 1 µM MPA. Experiments were repeated three times. L. 3D-projection of images of the nuclear Impdh2-cytoophidia (red) in macrophages after RANKL treatment. The Impdh2-cytoophidia structures were visualized by immunofluorescence. Scale bar, 3 µm. M. The ratio of Impdh2-cytoophidia in the cytoplasm and nucleus of the macrophages after RANKL treatment. Eight random microscope fields of cells were counted. In (B, F-H, M), results were expressed as mean ± s.d. \*\**p*<0.01, \*\*\**p*<0.001, \*\*\*\**p*<0.0001, versus Wt, Student’s *t*-test. See also Supplementary Figure S7.

Similarly, we next investigated whether nuclear cytoophidia assembly suppresses Impdh2 transcriptional activity during osteoclastogenesis. The number of multinucleated osteoclasts was dramatically decreased after MPA treatment (Fig 7G). The expression of osteoclastic transcription factors and marker genes was dramatically decreased during osteoclastogenesis. MPA treatment slightly increased the Impdh2 protein expression level without overtly affecting the mRNA expression (Fig 7H and 7I). Furthermore, the OXPHOS complexes were suppressed by MPA treatment (Fig 7J). The expression of Impdh2 was increased in the nucleus (Fig 7K), which also appeared to contradict the results that Impdh2 deficiency inhibits osteoclast differentiation. Furthermore, we observed that the nuclear Impdh2-cytoophidia existed in osteoclasts *in vivo* (Supplementary Fig S7C), and were significantly accumulated by MPA treatment *in vitro* (Fig 7L, 7M, and Supplementary Fig S7D), indicating that Impdh2-cytoophidia assembly also suppressed the transcriptional activity of Impdh2 during osteoclast differentiation. To further evaluate the function of the Impdh2-cytoophidia assembly or disassembly on osteoclast differentiation, macrophages were washed by PBS after MPA treatment for 24 h and then were stimulated with RANKL on the next day (Supplementary Fig S7E). Without MPA treatment, the canonical Impdh2- cytoophidia structures were only detected in a few macrophages (Supplementary Fig S7D). A large number of the nuclear Impdh2-cytoophidia were assembled after MPA treatment and meanwhile, inhibited the osteoclasts formation. Intriguingly, once MPA was deprived, the number of osteoclasts was recovered to comparable levels with the control group. Furthermore, the Impdh2-cytoophidia disassembled and osteoclast differentiation recovered (Supplementary Fig S7F and S7G), suggesting that cytoophidia assembly and disassembly are dynamic and reversible. Collectively, these results confirmed that nuclear cytoophidia assembly suppresses Impdh2 transcriptional activity.

## DISCUSSION

Compartmentalization via filamentation is a class of regulatory mechanisms for biological processes through protein condensation or protein-protein interactions (Lynch, Kollman et al., 2020). Cytoophidia formation regulates the inherent activity of proteins. However, a key question is whether the assembled cytoophidia promote or suppress the physiological function of these proteins, which has not yet been resolved. In this study, we show that the transcriptional regulation of Impdh2 is required for limb formation and bone resorption, and nuclear filamentation inhibits the transcriptional activity of Impdh2.

Cytoophidium has recently been discovered in many species and is a new research frontier in the cell biology field. So far, about 23 proteins, mainly composed of metabolic enzymes, have been identified to be able to assemble into cytoophidia, including cytidine-5’-triphosphate synthase (Ctps) and asparagine synthetase (Asns) (Shen, Kassim et al., 2016). The mutation or deficiency of these proteins assembling the cytoophidia will cause severe diseases, such as CTPS1 deficient patients cause a novel and life-threatening immunodeficiency (Martin, Minet et al., 2020, Martin, Palmic et al., 2014); ASNS deficiency results in a rare neurometabolic disease manifesting as microcephaly, severe developmental delay, and spastic quadriplegia (Liu, Wang et al., 2022, Ruzzo, Capo-Chichi et al., 2013); IMPDH1 mutation cause blindness (Aherne *et al*, 2004, Burrell *et al*, 2022). Interestingly, these proteins, similar to Impdh2, also exist in the nucleus. Thus, we propose that these cytoophidia-assembly proteins might also be transcription factors to regulate the physiopathological processes.

Eukaryotic cytoophidia encompass cytoplasmic cytoophidia and nuclear cytoophidia. Due to the highly abundant cytoophidia in the cytoplasm, almost all current research on the function of cytoophidia is focused on the cytoplasmic cytoophidia. Nuclear cytoophidia have similar components to the cytoplasmic cytoophidia. Therefore, they might share some similar functions with the cytoplasmic cytoophidia, such as prolonging the half-life of proteins (Chang *et al*, 2022, Sun & Liu, 2019). Nevertheless, because of the difference in spatial location, nuclear cytoophidia might have distinct functions from the cytoplasmic cytoophidia. IMPDH1-assembled cytoophidia translocate the metastasis-related gene Y-box binding protein 1 (YB-1) into the nucleus via physical interaction to regulate the metastasis in clear cell renal cell carcinoma (Ruan *et al*, 2020). In this study, we demonstrated that nuclear filamentation inhibits the transcriptional activity of Impdh2 during bone development and resorption.

Mesenchymal stem cells are a heterogeneous subset of stromal stem cells that can differentiate into various types of cells, including adipocytes, osteoblasts, osteocytes and chondrocytes, as well as cells of other embryonic lineages (Uccelli *et al*, 2008). Impdh2, required for life, ubiquitously expresses in most tissues (Gu, Stegmann et al., 2000, Senda & Natsumeda, 1994). Furthermore, Impdh2-cytoophidia naturally form in embryonic stem cells (Carcamo *et al*, 2014). In the current study, we confirmed that Impdh2 functions in the limb mesenchymal progenitors and is essential for bone development. Thus, we speculate that Impdh2 also functions, potentially through cytoophidia, in other cell lineages derived from mesenchymal stem cells, e.g., adipocytes and osteoblasts. On the other hand, cytoplasmic Impdh-cytoophidia mediate HIF-1α signaling pathway regulation in disuse osteoporosis (Bie *et al*, 2023), and our study found that deletion of Impdh2 inhibits osteoclastogenesis via the impaired function of the mitochondrion and nuclear Impdh2-cytoophidia repress its transcriptional activity, indicating that Impdh2 regulates bone resorption by multifaceted regulatory mechanisms.

Cytoophidia structures could provide a platform to regulate the function of other proteins by direct physical interactions. To date, several proteins have been identified to favor the cytoophidia assembly through interaction with Impdh2, for instance, ARL2 (Schiavon, Griffin et al., 2018), CTPS (Chang *et al*, 2018, Keppeke *et al*, 2015), and ANKRD9 (Hayward *et al*, 2019). Thus, what the roles of these interaction proteins are and whether they function via Impdh2-cytoophidia should be addressed in the future. In summary, we genetically demonstrated a previously unrecognized function of Impdh2 on bone development and resorption, and identified, for the first time, the nuclear cytoophidia assembly inhibits the transcriptional activity of impdh2. The genetic mouse models and the new regulatory mechanism by nuclear cytoophidia assembly will be instrumental for future studies to further explore the functions of Impdh2 on skeletal development and homeostasis.

### Limitations of the study

Impdh has been demonstrated to be a transcription factor to regulate cell proliferation in *Drosophila* (Kozhevnikova, van der Knaap et al., 2012). In this study, we showed that Impdh2 transcriptional regulates skeletal development and homeostasis in mice. However, given the evolutionary conservation of Impdh2, we do not investigate whether Impdh2 also functions as a transcription factor in other tissues or species. In addition, although we demonstrated that nuclear filamentation inhibits the transcriptional activity of Impdh2 during limb formation and bone resorption, evidence has shown that IMPDH2 are associated with neurodevelopmental disorders (Lake *et al*, 2016, O’Neill *et al*, 2023) and inflammation-relevant diseases (Liao *et al*, 2017). This evidence does not rule out a possible role for the nuclear Impdh2-assembled cytoophidia function in these pathological processes.

## MATERIALS AND METHODS

### Primary cell cultures

For micromass culture, limb buds from the E12.5 mouse embryo were isolated and digested with 1 mg/ml type II collagenase (Worthington, LS004176) for 1 h at 37 °C. The cells were filtered with the 40 μm cell strainer, centrifuged at 1,000 rpm for 5 min and resuspended in DMEM:F-12 (1:1) (Gibco, C11330500BT) media supplemented with 10% FBS and 1% penicillin- streptomycin. After reconstituting at a density of 1×10^7^ cells/ml, 10 μl of the mesenchymal cells (1×10^5^ cells in total) were dropped into the 24-well plate and incubated for 1 h in 37°C incubator for attachment. The cells were then added the fresh DMEM:F-12 (1:1) media supplemented with 10% FBS, 1% penicillin-streptomycin, 50 mg/ml ascorbic acid (Sigma-Aldrich, A4544) and 10 mM β-glycerol phosphate (Sigma-Aldrich, G9422) for 6 days until the chondrogenic nodule formation. The cells were harvested for alcian blue staining, western blot and qPCR analysis.

For osteoclast differentiation, mouse bone marrow cells from age- and gender-matched wild or mutant mice were harvested and cultured for 3 days in α-MEM medium with 10% FBS, 1% penicillin-streptomycin, 1% L-glutamine, and L929 cell supernatant (conditioned medium, herein referred as CM), which contained the equivalent of 20 ng/ml of murine rM-CSF and was used as a source of M-CSF. The attached bone marrow–derived macrophages (BMMs) were scraped, and 4.5×10^4^ cells/cm^2^ were seeded in α-MEM medium with 10% FBS, 1% penicillin-streptomycin, 1% L-glutamine, and CM for overnight, and then stimulated by 40 ng/ml RNAKL (PeproTech, 310-01) and CM. The media was changed after 3 days. The differentiated osteoclasts were evaluated by TRAP staining.

### Mouse models

Floxed *Impdh2* mice, *Rosa26*-*ZsGreen* mice, and transgenic mice expressing *Cre*-recombinase under the control of the *Prx1* promoter and lysozyme M promoter were purchased from Jackson Laboratory. *Prx1*-*Cre* or *LysM*-*Cre* mice were mated with *Impdh2^f/+^* mice to generate the limb mesenchymal cells or myeloid lineage-specific knockout mice (hereafter referred to as *Impdh2^Prx1-/-^*or *Impdh2^LysM-/-^*), respectively. We used the littermates *Impdh2^f/f^*mice (hereafter referred to as Wt) as the control in the experiments. All mice were maintained under standard 12 h light/dark cycles with ad libitum access to regular food and water. All animal studies were performed with the approval of the Medical Ethics Committee of Jianghan University and conformed to relevant guidelines and laws. All mouse lines were kept on a C57BL/6 background.

For OVX experiments, bilateral ovariectomy or sham operation was performed on 10-week-old female mice from Wt and *Impdh2^LysM-**/**-^*mice. Mice were sacrificed 5 weeks after surgery. The uterus was carefully dissected and weighed for each mouse to confirm the success of the OVX operation. The femurs were collected for micro–computed tomography (µCT) analysis.

### Mouse skeletal preparation

The neonates were stained with alcian blue and alizarin red as previously described (Takarada *et al*, 2013). In brief, the pups were eviscerated, and the skins were carefully removed. The neonates were fixed in 95% ethanol overnight, then immersed in the alcian blue solution (0.015% alcian blue 8GX in 20% acetic acid and 80% ethanol) for overnight. The specimens were transferred to 2% KOH for 24 h after rinsing with 95% ethanol at least 3 h, and subsequently stained in alizarin red solution (0.005% alizarin sodium sulfate in 1% KOH) for overnight. Finally, the skeletons were cleared in 1% KOH/20% glycerol for at least 2 days and stored in 50% ethanol/50% glycerol.

### Whole-mount *in situ* hybridization

Whole-mount in situ hybridization and the digoxygenin-labeled riboprobes were conducted according to the published protocol (Rutkowsky *et al*, 2014). Briefly, after fixation with 4% paraformaldehyde (PFA) overnight, embryos were dehydrated through a methanol gradient (25, 50, 75, and 2×100% methanol in PBST) at room temperature. The specimens were bleached by 6% H_2_O_2_ for 1 h, incubated in 10 μg/mL proteinase K (Sangon Biotech, B600169- 0600) in PBST for 15-20 min depending on embryo stages, and immersed in the hybridization buffer (50% deionized formamide, 5×SSC, PH 4.5, 1% SDS, 50 μg/ml yeast tRNA, 50 μg/mL heparin) containing the appropriate probes for 20-24 h at 70 °C. Then, the specimens were sequentially washed several times through solution I (50% Formamide, 5×SSC pH 4.5, 1% SDS), solution II (0.5 M NaCl, 10 mM Tris–HCl pH 7.5, 0.1% Tween-20) and solution III (50% Formamide, 2×SSC pH 4.5, 0.1% Tween-20). After incubation with the blocking solution (10% sheep serum, 0.2% BMB blocking reagent in 1×TBST) for 1.5 h at room temperature, embryos were incubated with anti-DIG-AP Fab fragments antibody (Roche, 11093274910, 1:2000) for overnight. The specimens were washed by TBST 5 times for 1 h each, and developed with BM Purple AP substrate (Roche, 11442074001). Embryos were then refixed with 4% PFA and rinsed by PBST. Finally, they were visualized and imaged using a stereoscope and digital camera. The antisense riboprobes were synthesized with DIG RNA Labeling Mixture (Sigma-Aldrich, 11277073910) by PCR amplification using the following primers: *Impdh2*, forward 5’- GTCCTTAGCCCCAAGGATCG-3’, reverse 5’- GCGCGATAATACGACTCACTATAGGGCCACAGGCCAACACTTCCT-3’; *Sox9*, forward 5’- GAAGTCGCTGAAGAACGGACAAG-3’, reverse 5’- GCGCGATAATACGACTCACTATAGGGGCTGTAGTGAGGAAGGTTGAAGG G-3’; *Col2a1*, forward 5’- CCAGGTGAAGGTGGAAAGCAAG-3’, reverse 5’- GCGCGATAATACGACTCACTATAGGGTCAAGACCAGAGGGACCATCATC-3’; *Hoxd13*, forward 5’- TGGGCTATGGCTACCACTTC-3’, reverse 5’- GCGCGATAATACGACTCACTATAGGGTGTCCTTCACCCTTCGATTC-3’.

### Western blot

The whole cell protein extracts were collected using lysis buffer containing 150 mM Tris-HCl (pH 6.8), 6% SDS, 30% glycerol, and 0.03% bromophenol blue. 10% 2-mercaptoethanol was added before harvesting the cells. Cell lysates were separated by SDS-PAGE, transferred to Immobilon-P membranes (Millipore, Sigma), incubated with the primary antibodies and developed with a horseradish peroxidase-conjugated goat anti-mouse (BioRad, 1706516, 1:5,000), anti-rabbit (BioRad, 1706515, 1:5,000) or anti-rat (Jackson ImmunoResearch, 112-035-003, 1:5,000) immunoglobulin (IgG) antibodies, and detected using Clarity Western ECL Substrate detection kits (BioRad, 1705061). Mouse anti-Nfatc1 (556602, 1:1,000) was purchased from BD Biosciences; rat monoclonal anti-Blimp1 (sc-47732, 1:1,000) and mouse monoclonal anti-c-Fos (E-8) (sc-166940, 1:1,000) were purchased from Santa Cruz Biotechnology; rabbit polyclonal anti-Impdh2 (12948-1-AP, 1:1,000) and mouse monoclonal anti-Lamin B1 (66095-1, 1:20,000) were purchased from Proteintech; rabbit monoclonal anti-Sox9 (A19710, 1:1,000) and mouse monoclonal anti-Gapdh (AC002, 1:20,000) were purchased from ABclonal; Total oxidative phosphorylation (OXPHOS) Rodent WB Antibody Cocktail (ab110413, 1:1,000) was purchased from Abcam.

### Reverse transcription and real-time PCR

Total RNAs were extracted using TRIzol reagents and 0.5 μg of RNA was reverse transcribed using a HiScript Q RT SuperMix qPCR (+gDNA wiper) Kit (Vazyme, R123-01) according to the manufacturer’s instructions. Quantitative analysis of gene expression was performed in triplicate using the Applied Biosystems Real-time PCR System and ChamQ SYBR qPCR Master Mix kit (Vazyme, Q311-02). Gapdh expression was used as an internal control. The list of all primers used in this study was shown in Supplementary Table S1.

### Bone phenotype analysis

For primary ossification center (POC) and chondrocyte analysis in *Impdh2^Prx-/-^* mice, the femur, humerus and sternum of neonates (P0) were collected and then fixed with 4% PFA overnight. The bones were decalcified with 14% EDTA at 4 °C for 10 days. All samples were embedded in paraffin and sliced into 5 µm thick sections. Sections were deparaffinized and hydrated. The bone sections were subsequently performed alcian blue staining and hematoxylin- eosin staining using the standard protocols.

For µCT analysis, the femora from 12-week-old male mice were fixed in 4% PFA and scanned at 6 µm resolution using the µCT scanner (Bruker, SkyScan 1276). Distal femoral trabecular bone parameters or the cortical bone parameters obtained from the midshaft of femurs were analyzed using CT-Analyser (CTAn) software according to the manufacturer’s instructions and the American Society of Bone and Mineral Research (ASBMR) guidelines.

For bone histomorphometric analysis, 12-week-old male mice were intraperitoneally injected with calcein (25 mg/kg; Sigma, C0875) at 5 days and 2 days before euthanasia to obtain double labeling of newly formed bones. The third and fourth undecalcified lumbar vertebrae (L3 and L4) were embedded in methyl methacrylate (MMA)-based resin. 5 μm thick sections were cut using a Leica RM2255 rotary microtome (Leica Microsystems). Undecalcified sections were performed with von Kossa, tartrate-resistant acid phosphatase (TRAP) or toluidine blue staining. Osteomeasure software (OsteoMetrics) was used for static and dynamic histomorphometric analysis following standard procedures according to the manufacturer’s instructions previously described (Xu *et al*, 2016).

### Immunofluorescence staining

For the cultured limb mesenchymal cells or osteoclasts staining, cells were fixed with 4% PFA for 10 min, washed three times with PBS, and permeabilized with 0.2% Triton X-100 for 15 min at room temperature. The cells were blocked with 1% BSA/2.5% donkey serum for 1 h and incubated with primary antibodies at 4 °C for overnight. Alexa Fluor™ 568 Phalloidin (1:500) was used for labeling the F-actin of osteoclasts. For tissue staining, after fixation by 4% PFA, the mouse decalcified femur treated with 14% ethylenediaminetetraacetic acid (EDTA) for 7 d, and forelimbs from E12.5 embryo were equilibrated in 30% sucrose for overnight before embedding in OCT media. Cryosections were collected at 5 to 7 μm, blocked with 5% donkey serum (Sigma-Aldrich, D9663) for 2 h, and then incubated with primary antibodies at 4 °C overnight. Mouse monoclonal anti-cathepsin K (Santa Cruz Biotechnology, sc-48353, 1:200) was used to label the osteoclasts of bone. Rabbit polyclonal anti-Impdh2 (1:1,000) was used to detect the Impdh2-assembled cytoophidia. All samples were incubated with the secondary antibody conjugated with donkey anti-rabbit Alexa Fluor Plus 488 (Invitrogen, A32790, 1:500) and donkey anti-mouse Alexa Fluor Plus 594 (Invitrogen, A32744,1:500) for 1 h at room temperature. Nuclei were stained with hoechst for 5 min (1:2,000) after washing with PBS one time. Samples were then washed with PBS and mounted with Fluoromount-G. All fluorescent images were obtained on the Leica SP8 confocal microscope.

### RNA sequencing and analysis

Total RNAs were extracted using TRIzol reagents following the manufacturer’s instructions. Hieff NGS™ MaxUp Dual-mode mRNA Library Prep Kit for Illumina® (YEASEN, 12301ES96) was used to purify poly-AC+Ctranscripts and generate sequencing libraries following the manufacturer’s instructions. High-throughput transcriptome sequencing of RNAs (RNA-seq) was performed at Shanghai Shenggong Biotech Co., Ltd as described.(Cheng *et al*, 2021) RNA-seq reads were aligned to the mouse genome assembly (GRCm39/mm39) using HISAT2 (version 2.1.0) with default parameters.(Kim *et al*, 2015) Fragments Per Kilobase of exon model per Million mapped fragments (FPKM) was calculated by StringTie (version 1.3.3b) to quantify the transcript abundances. DEGseq (version 1.26.0) was used for differential gene expression analysis. To obtain the significant differential genes regulated by Impdh2 deletion in limb mesenchymal cells, the screening conditions were set as follows: |log2FoldChange|>0.5 and *p*-value<0.05. To obtain the significant differential genes regulated by Impdh2 deletion in osteoclastogenesis, the screening conditions were set as follows: |log2FoldChange|>0.3 and *p*-value<0.05. Gene Ontology (GO) analysis was performed using the Database for Annotation, Visualization, and Integrated Discovery (DAVID) Functional Annotation tool. Impdh2-regulated pathways by the GO analysis were ranked based on the *p* values.

### Statistical analysis

Statistical analysis was performed using GraphPad Prism *version 8.0* software. Two-tailed Student’s *t*-test was applied to test for differences between two groups of data. *p*<0.05 was taken as statistically significant. All data are presented as the mean ± s.d. as indicated in the figure legends. All experiments were repeated at least three times.

## DATA AVAILABILITY

Bulk RNA-seq data have been deposited in the Gene Expression Omnibus database with the accession code GSE244054.

## ACKNOWLEDGMENTS

We thank Rongxiang He for his help to obtain the micrographs by scanning electron microscope, Gang Cao for methyl methacrylate (MMA)-based resin samples preparation. The team was funded by the National Key R&D Program of China (grant number: 2020YFA0907400), the National Natural Science Foundation of China (grant number: 32170702 and 82000828), the Natural Science Foundation of Hubei Province (2022CFB281), the Wuhan Science and Technology Bureau of Hubei Province of China (20222508770001) and the Major Special Funding Program for First-class Discipline Construction of Jianghan University (grant number: 2023XKZ021).

## AUTHOR CONTRIBUTIONS

C.X. and Z.H. designed experiments; C.X., Z.W., L.L., X.D., W.X., J.X., H.L., X.Z., Y.L., W.W., H.J., Y.G., C.L., K.X., S.W. and Y.A. performed experiments; C.X., Z.W., L.L., X.D. and Z.H. analyzed data; C.X. and Z.H. supervised the project and wrote the manuscript.

---

**Supplementary Figure S1.**
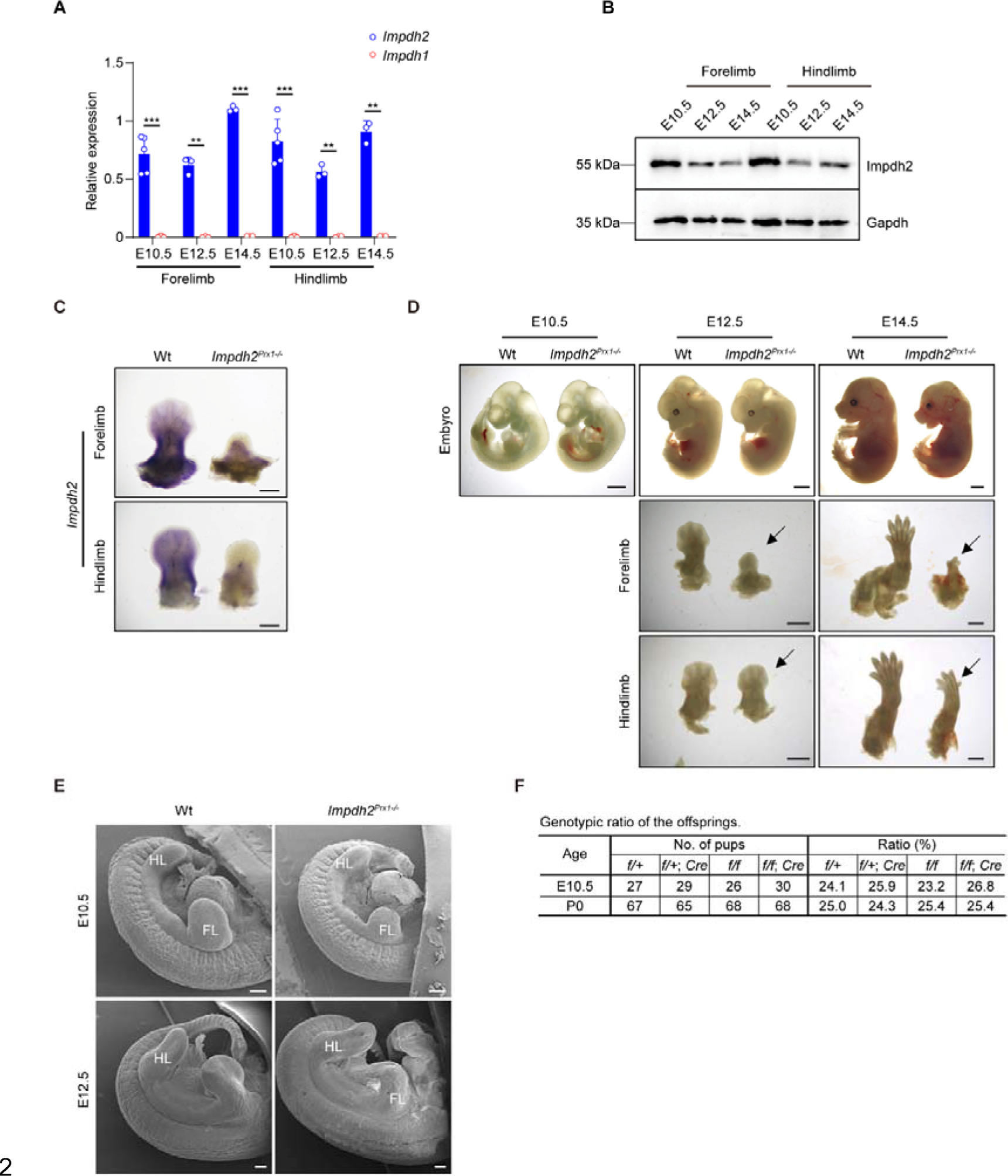
Deletion of Impdh2 induces abnormalities during limb development. A. Quantitative qPCR analysis of *Impdh1* (blue) and *Impdh2* (red) expression in the forelimbs and hindlimbs. Total RNAs were extracted from the forelimbs and hindlimbs of Wt embryos at the indicated stages. Note that compared with *Impdh1*, the expression of *Impdh2* was much higher in the limb buds. E10.5, n=5; E12.5 and E14.5, n=3. B. Western blot analysis of Impdh2 expression in the forelimbs and hindlimbs extracted from mouse embryos at the indicated stages. Experiments were repeated three times. C. Whole-mount *in situ* hybridization for *Impdh2* in the forelimbs and hindlimbs from Wt and *Impdh2^Prx1-/-^* embryos at E12.5. Experiments were repeated three times. Scale bar, 400 µm. D. Gross appearance of the whole embryos, forelimbs and hindlimbs from Wt and *Impdh2^Prx1-/-^* mice at the indicated stages. Note that limbs of *Impdh2^Prx1-/-^* embryo showed abnormal development (arrow). Scale bar, 1 mm. E. Micrographs of limb buds from Wt and *Impdh2^Prx1-/-^* embryos by scanning electron microscope at E10.5 and E12.5. Scale bar, 200 µm. F. The different genotype ratios of offsprings from female *Impdh2^f/f^*; +/+ crossed with male *Impdh2^f/f^*; *Prx-Cre/+*. In (A), results were expressed as mean ± s.d. \*\**p*<0.01, \*\*\**p*<0.001, versus Wt, Student’s *t*-test. Related to Figure 1.

**Supplementary Figure S2.**
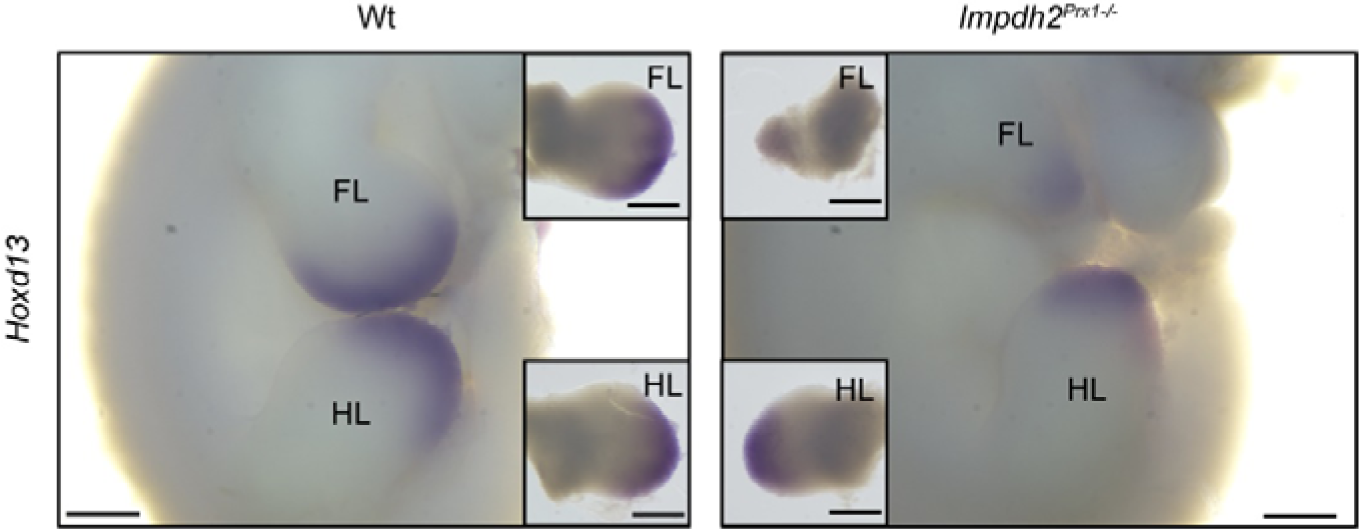
Deletion of Impdh2 impairs *Hoxd13* gene expression. Whole-mount *in situ* hybridization for *Hoxd13* from Wt (left) and *Impdh2^Prx1-/-^* (right) embryos at E11.5. FL: forelimb, HL: hindlimb. Scale bar, 400 µm. Related to Figure 3.

**Supplementary Figure S3.**
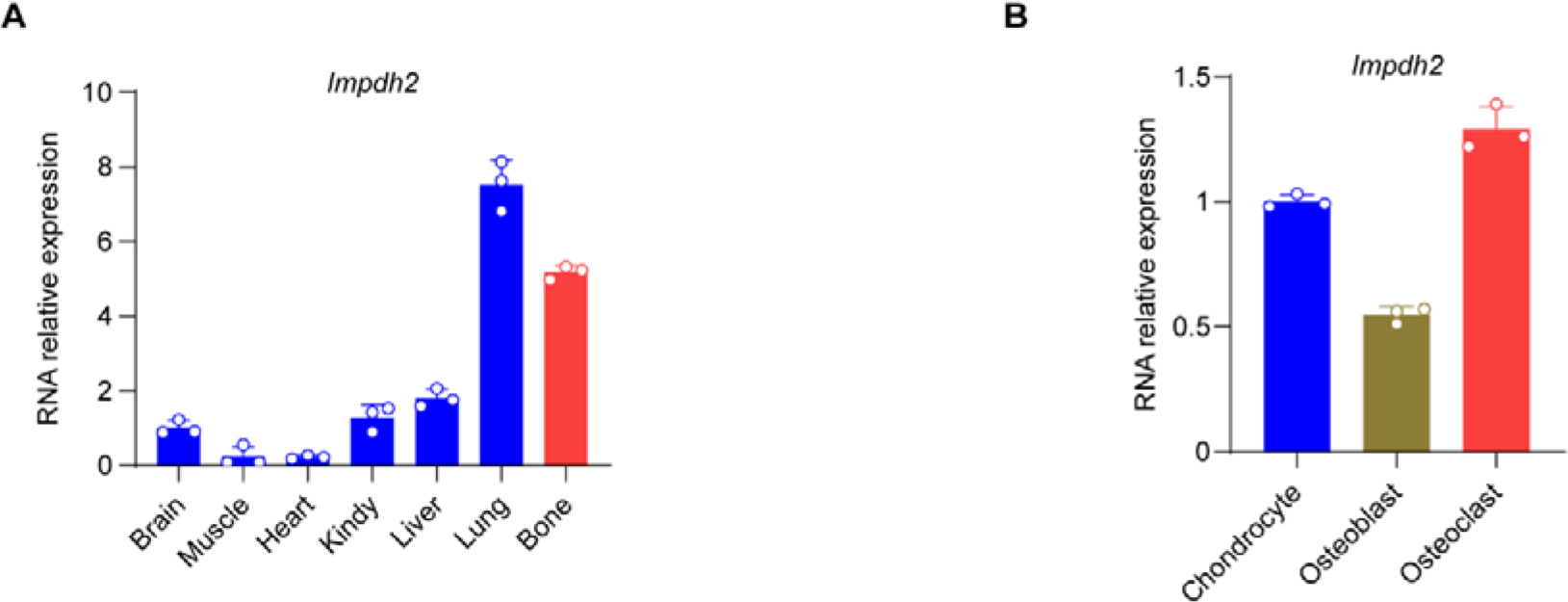
*Impdh2* expresses in different tissues and bone-related cell lineages. A. Quantitative qPCR analysis of *Impdh2* mRNA expression in mouse primary tissues. Experiments were repeated three times. B. Quantitative qPCR analysis of *Impdh2* mRNA expression in mouse primary chondrocytes, osteoclasts, and osteoclasts. Experiments were repeated three times. Related to Figure 5.

**Supplementary Figure S4.**
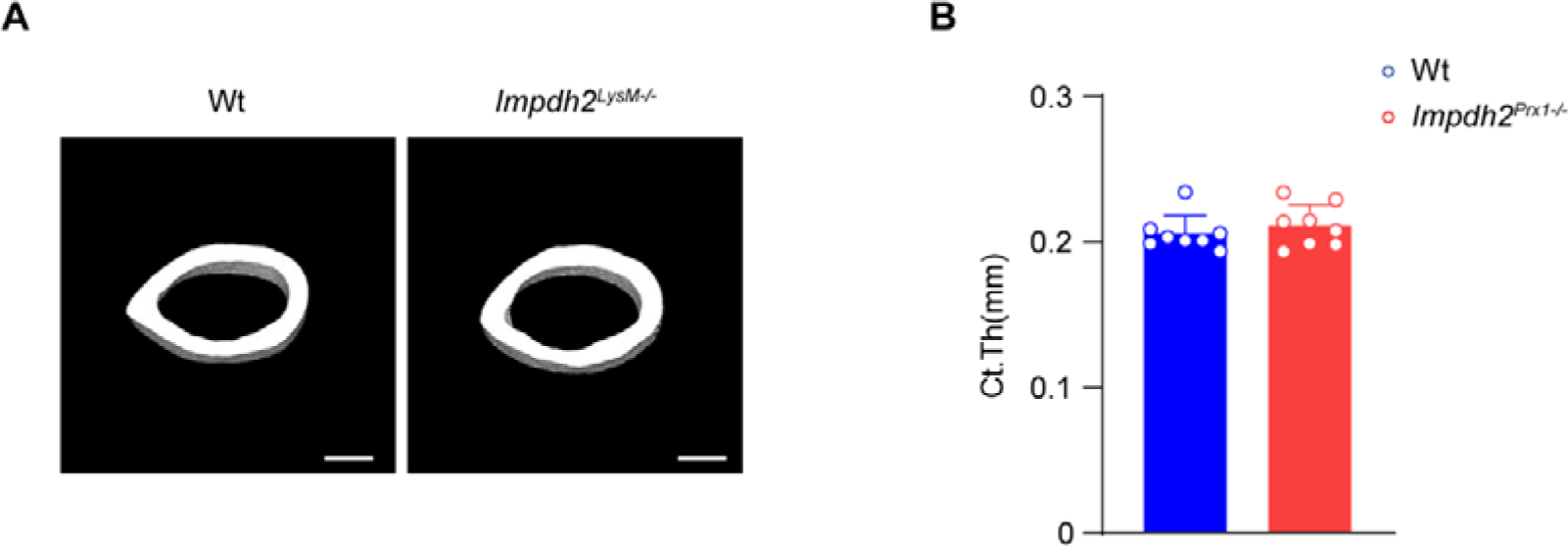
Impdh2 deletion does not affect cortical thickness. A, B. µCT images (A) and bone morphometric analysis (B) of cortical bone of the mid-shaft femurs isolated from 12-week-old male Wt and *Impdh2^LysM-/-^*mice. n=8/group. Versus Wt, Student’s *t*-test. Results were expressed as mean ± s.d. Scale bar, 500 µm. Related to Figure 5.

**Supplementary Figure S5.**
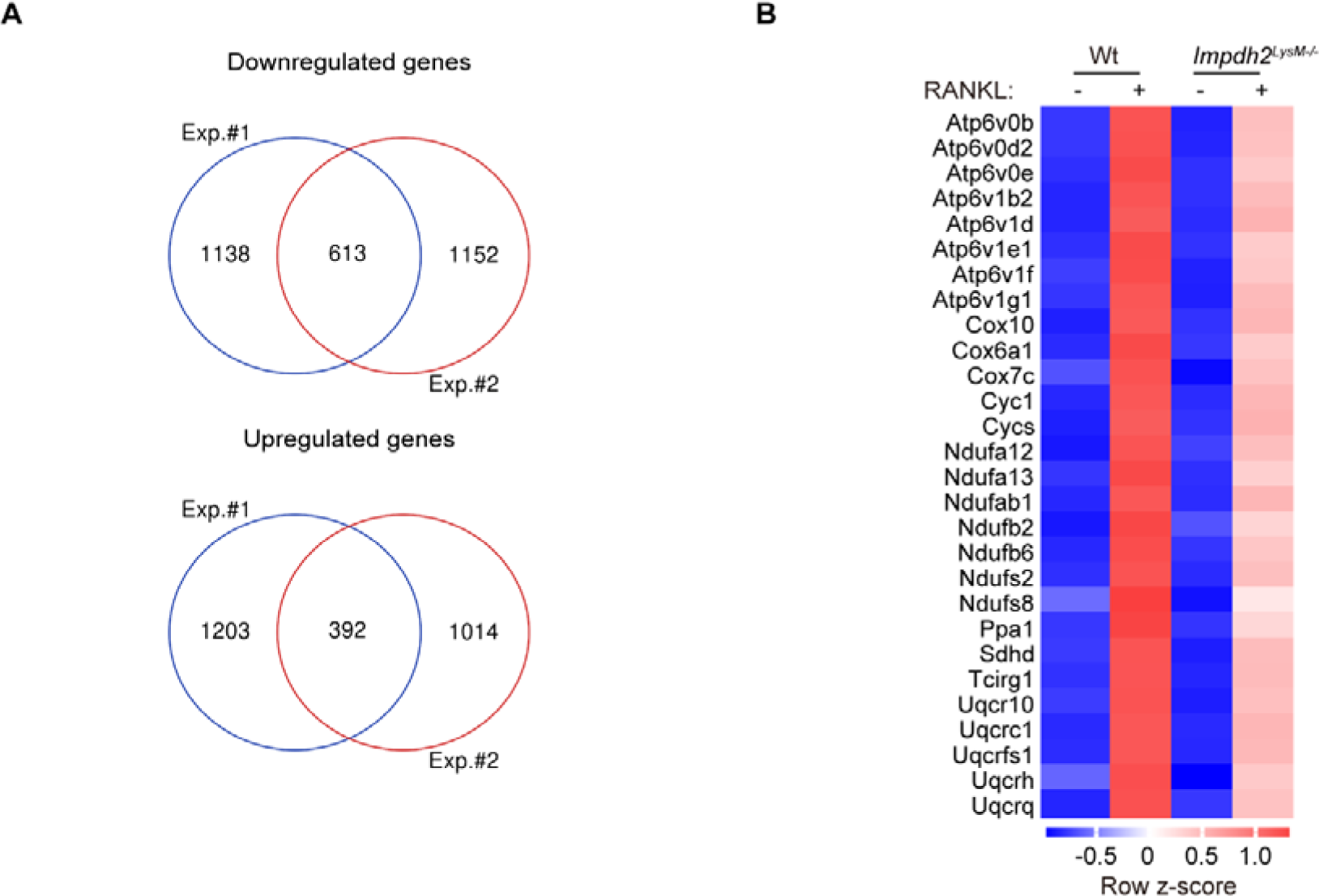
Mitochondrial OXPHOS expression is impaired by RNA-seq analysis. A. Venn diagrams of downregulated or upregulated genes by Impdh2 deficiency from 2 batches of osteoclast differentiation experiment. Exp. #1, experiment #1; Exp. #2, experiment #2. B. Heatmaps of mRNA expression of OXPHOS-related downregulated genes by Impdh2 deficiency. Row z-score of FPKMs of genes was shown in the heatmap. Related to Figure 6.

**Supplementary Figure S6.**
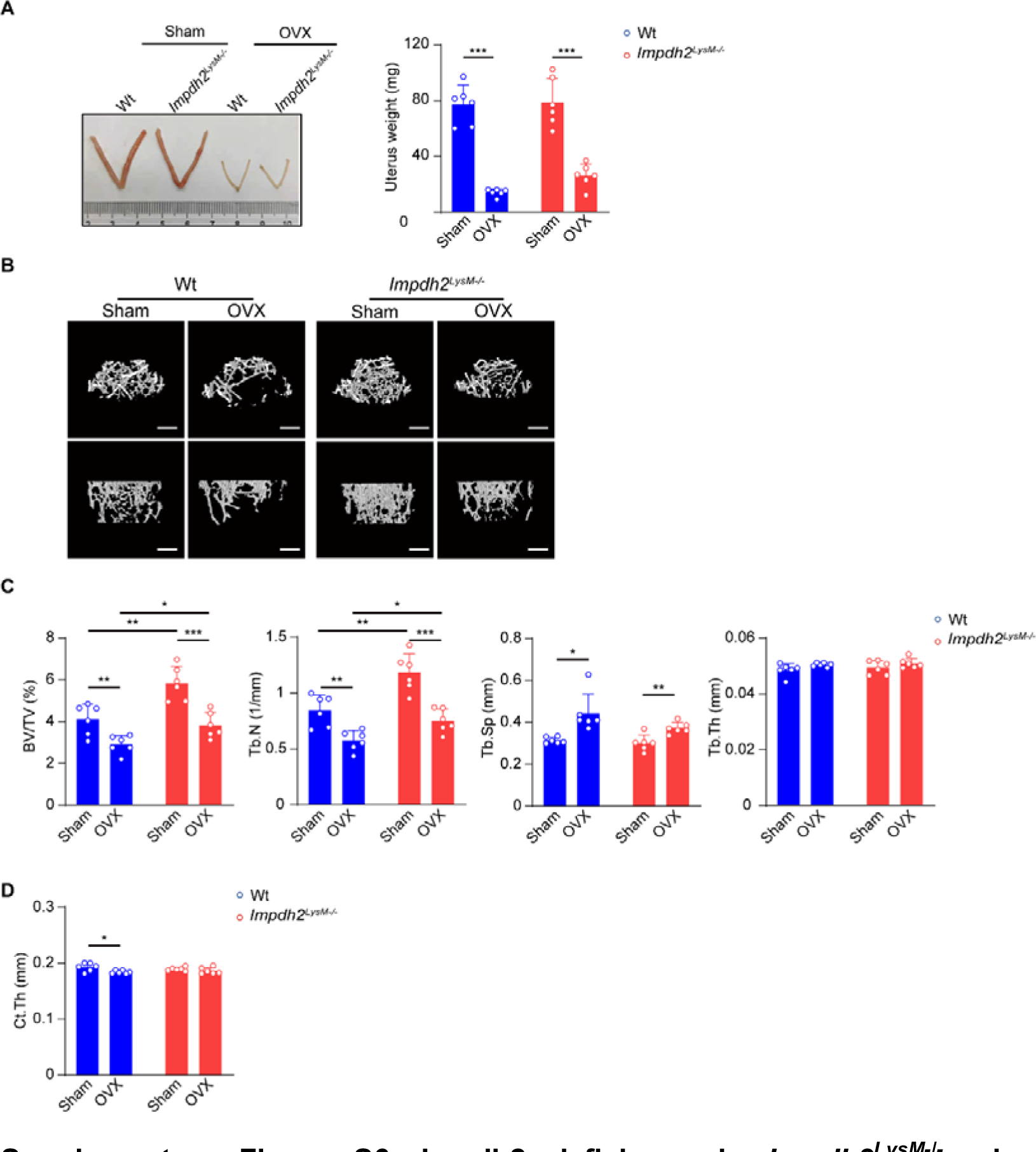
Impdh2 deficiency in *Impdh2^LysM-/-^* mice prevents OVX-induced osteoporosis. A. Representative image (left) and weight (right) of the uterus. n=6/group. B, C. μCT images (B), and bone morphometric analysis (C) of trabecular bone of the distal femurs isolated from the Wt and *Impdh2^LysM-/-^* female mice with sham or OVX surgery. BV/TV, bone volume/tissue volume; Tb.N, trabecular number; Tb.Sp, trabecular separation; Tb.Th, trabecular thickness; Ct.Th, cortical thickness. n=6/group. Scale bar, 500 µm. D. Cortical thickness (Ct.Th) analysis of cortical bone of the mid-shaft femurs isolated from the Wt and *Impdh2^LysM-/-^* female mice with sham or OVX surgery. n=6/group. In (A, C, D), results were expressed as mean ± s.d. \**p*<0.05, \*\**p*<0.01, \*\*\**p*<0.001, versus Wt, Student’s *t*-test. Related to Figure 6.

**Supplementary Figure S7.**
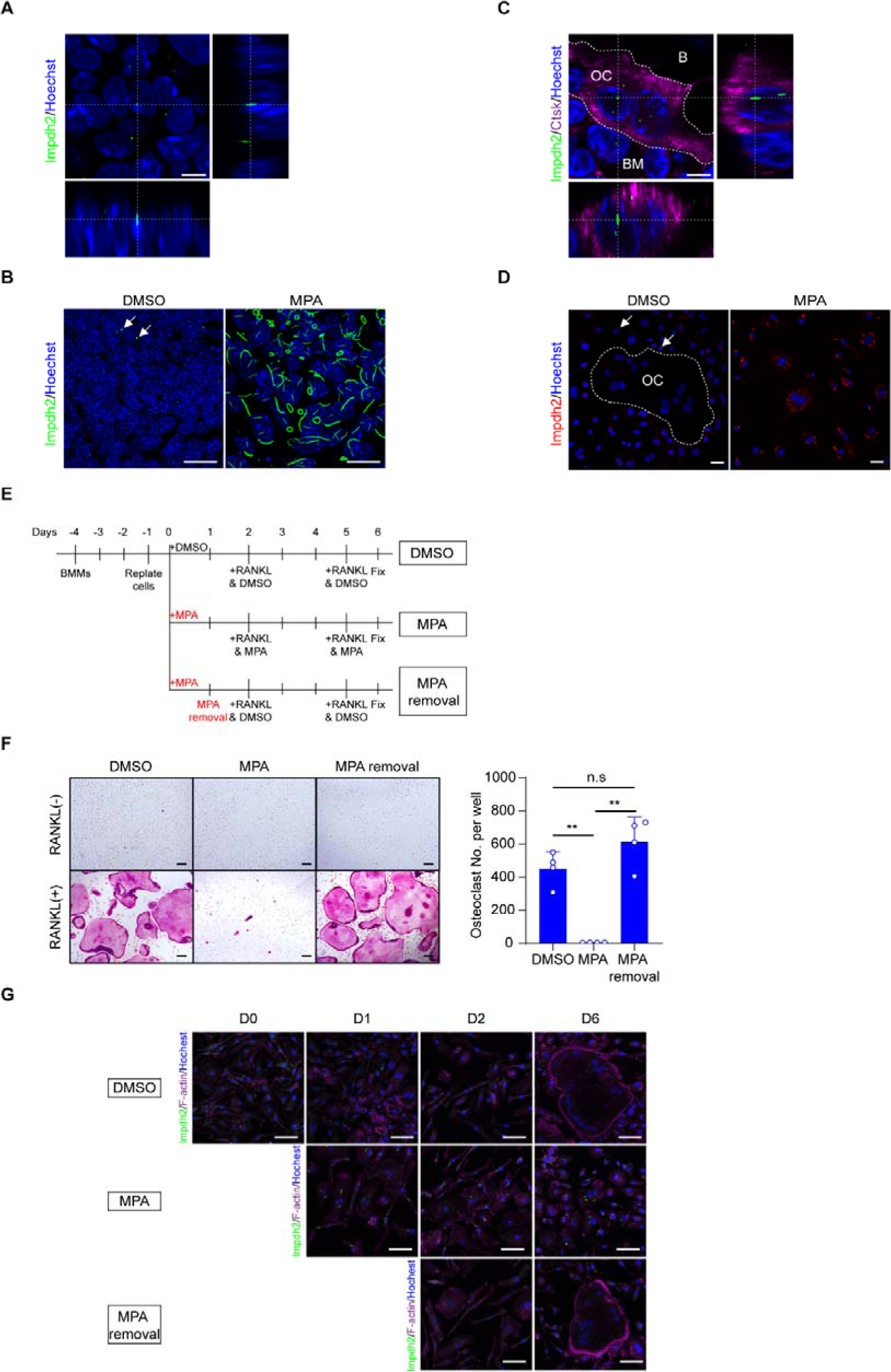
Impdh2-cytoophidia are induced by MPA treatment. A. 3D-projection of images of the nuclear Impdh2-cytoophidia (green) in Wt forelimbs at E10.5. Scale bar, 4 µm. B. Immunofluorescence staining images of Impdh2-cytoophidia (green) in primary mesenchymal progenitor cells isolated from Wt limbs at E12.5 treated with DMSO (left) or 1 µM MPA (right). Impdh2-cytoophidia were formed under the normal condition (arrows). Scale bar, 20 µm. C. 3D-projection of images of the nuclear Impdh2-cytoophidia (green) in the osteoclast of bone. Osteoclasts were marked by anti-Ctsk antibody (magenta). B, bone. OC, osteoclast. BM, bone marrow. The osteoclast was labeled as the dashed line. Scale bar, 4 µm. D. Immunofluorescence staining images of Impdh2-cytoophidia (red) in the macrophages stimulated with RANKL and treated with DMSO (left) or 1 µM MPA (right). Impdh2-cytoophidia were formed under the normal condition (arrows). OC, osteoclast. The osteoclast was marked as the dashed line. Scale bar, 20 µm. E. Schematic of experimental design of the cell culture for the following experiment. The macrophages derived from BMMs were replated and treated with DMSO or 1 µM MPA for 2 days, and then stimulated with RANKL for 4 days. For the MPA removal group, the macrophages were washed by PBS after 1 µM MPA treatment for 1 day, and after 24 h, stimulated with RANKL for 4 days. F. Osteoclast differentiation under the indicated conditions. TRAP staining was performed and the number of TRAP-positive multinuclear osteoclasts (≥3 nuclei/cell) per well was calculated. Scale bar, 100 µm. G. Representative images of immunostaining of Impdh2 (green) and F-actin (magenta) during osteoclast differentiation under the indicated conditions. Scale bar, 50 µm. In (F), results were expressed as mean ± s.d. \*\**p*<0.01, n.s., not statistically significant, versus Wt, Student’s *t*-test. Related to Figure 7.

**Supplementary Table S1.**
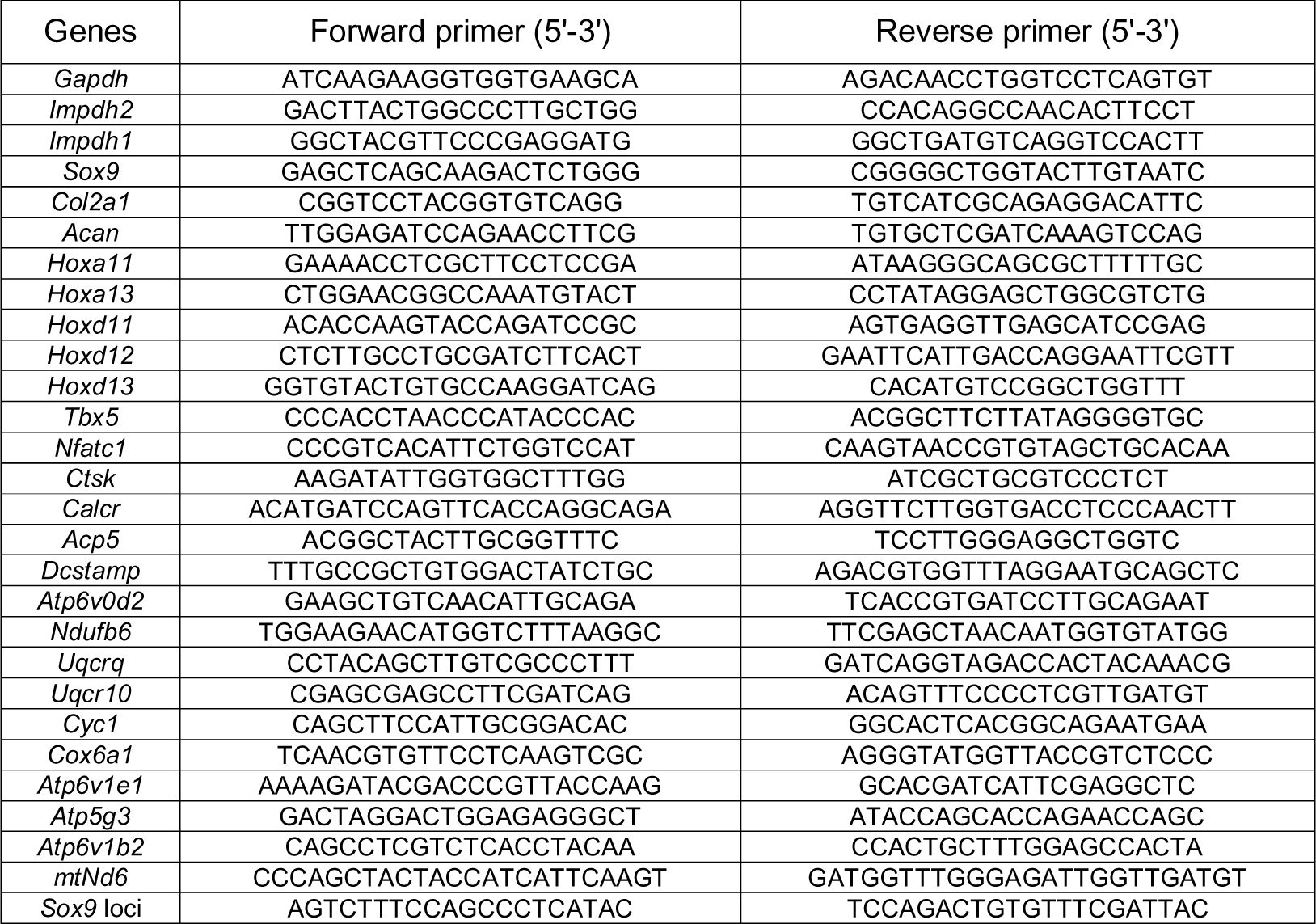
Oligonucleotide primers were used in this study.

